# CaPTrends: A global database of Carnivoran Population Trends

**DOI:** 10.1101/2022.01.13.476193

**Authors:** Thomas F. Johnson, Paula Cruz, Nick J. B. Isaac, Agustin Paviolo, Manuela González-Suárez

## Abstract

**Motivation:** Population trend information is an ‘Essential Biodiversity Variable’ for monitoring change in biodiversity over time. Here, we present a global dataset of 1122 population trends describing changes in abundance over time in large mammals from the Order Carnivora – some of the world’s most charismatic and functionally important fauna.

**Main types of variables included:** Key data fields for each record: species, coordinates, trend timeframe, methods of data collection and analysis, and population timeseries or summarised trend value. Population trend values are reported using quantitative metrics in 75% of records that collectively represent more than 6500 population estimates. The remaining records qualitatively describe population change (e.g. increase).

**Spatial location and grain:** Records represent locations across the globe (latitude: -51.0 to 80.0; longitude: -166.0 to 166.0) with more trends found within the northern temperate zone.

**Time period and grain:** Records span from 1726 to 2017, with 92% of trends starting after 1950.

**Major taxa and level of measurement:** We conducted a semi-systematic search for population trend data in 87 species from four families in the order Carnivora: Canidae, Felidae, Hyaenidae and Ursidae. We compiled data for 50 of the 87 species.

**Software format:** .csv

## Introduction

Rapid global change is threatening biodiversity (IPBES, 2019). However, biodiversity changes are not happening at the same rate in all places and species, with the fate of populations varying across regions (Fritz *et al*., 2009; Polaina *et al*., 2016), levels of protection (Amano *et al*., 2018), and the intrinsic traits of the affected species (Cardillo *et al*., 2005; Gonzalez-Suarez *et al*., 2013; González-Suárez & Revilla, 2013). An example of this variability in extinction can be seen in the largest terrestrial mammals in the order Carnivora, where there is evidence for population recoveries and recolonizations (Chapron *et al*., 2014), alongside declines and extinctions (Ripple *et al*., 2014).

Currently, the largest sources of mammalian population trend data are within BioTIME (Dornelas *et al*., 2018) and the Living Planet Index (WWF, 2020), which combined, provide millions of abundance observations. Here, we expand upon both datasets for four families in the order Carnivora: Canidae, Felidae, Hyaenidae and Ursidae - which represent some of the world’s most charismatic and functionally important fauna (Ripple *et al*., 2014). For the 87 species in these families, following the IUCN taxonomy, we compiled published population trend data from abundance time-series as in BioTIME and the Living Planet data (Dornelas *et al*., 2018; WWF, 2020). However crucially, we also searched for and included summarised estimates of change (e.g. mean population growth rate) and qualitative descriptions of population change, allowing the expansion of available data. These data provide the most comprehensive global overview of population status for these species and can be used to evaluate different factors that influence population changes and describe species’ status.

## Methods

### Locating population trend records

We used a systematic literature search to identify population trends in the primary literature. This search involved searching Scopus and Web of Science for population trend related terms (e.g. ‘population trend’, ‘declin*’ and ‘increas*’) alongside taxonomic information (e.g. species names). We searched for terms in English and Spanish. We found 30 articles in Spanish and 3233 articles in English. We narrowed down these articles to a highly relevant subset (i.e. likely to contain population trend information; N = 516) using titles and abstracts (see Supplementary: Systematic search). A selection of these highly relevant articles were syntheses of other studies – in this case, we referred to the primary source and included the article within our list, expanding the number of highly relevant articles to 536. We were unable to obtain the full text for 19 of these highly relevant articles, reducing our sample to 517 articles, which were to be read in full (see below).

### Extracting information from sources

When a source contained population trend information, we recorded the trend and additional metadata describing taxonomy, location, study period, and methodology (Table S1). Population changes were reported in a variety of formats, but broadly fall into two groups, quantitative where the trend was described numerically (e.g. %change), and qualitative where the trend was described categorically (e.g. increase). In the quantitative group, we record the trend as presented in the original source, and we recorded five distinct types: 1) population abundance time-series, 2) mean finite rate of population change (λ), 3) mean instantaneous rate of population change (*r*), 4) percentage change between two time points, and 5) fold change between two time points; further described in Table S1. For studies that reported trends in multiple formats, we recorded the most informative e.g. where raw abundance data were available this was preferred over summary estimates of population change. If the population values were only reported in a graph, we used a graphic digitiser (https://apps.automeris.io/wpd/) to estimate the values (Rohatgi, 2015).

For population trends calculated from time-series data, we recorded the length of the time-series (number of individual estimates used to derive the trend). For population trends based on matrix models and demographic parameters, we recorded the number of sampling years used to estimate the demographic parameters. For estimates of annual rates of change (λ and *r*) derived from three or more data points, we also noted any available estimate of dispersion (e.g. variance) and test-statistic values. For the qualitative descriptions of trends, we inferred the trend based on the description in the primary sources, with trends falling into the following four categories: increase – source described the population abundance as exhibiting overall growth during the monitored period; stable – source described the population abundance as exhibiting a stable or unchanged trend over the monitored period; decrease – source described the population abundance as exhibiting an overall decline during the monitored period; varied – source described the population abundance as exhibiting both growth and declines over the monitored period, without any clear directional trend. The specific terminology used to describe each trend varied between the primary sources, but the general message was largely consistent. However, we do acknowledge that each primary source likely has a different definition for a given trend (i.e. how much growth is necessary to be classed as an increase), which introduces an opportunity for inconsistency and subjectivity, and so these qualitative trends should be interpreted cautiously.

For each trend we recorded the binomial species name following the IUCN taxonomy – we report discrepancies between the IUCN taxonomy and another taxonomy (Wilson & Reeder, 2005) in Table S2. When the species name in the primary literature did not match the IUCN taxonomy, we referred to the list of IUCN taxonomy synonyms to locate the accepted IUCN species name. Subspecies names were also available in some primary sources, and we noted these as recorded in the primary source. For location, we recorded the name of the study site given in the primary source, whether the site was described as a protected area, and the country or countries it overlapped. If provided, we recorded the study site’s coordinates (minimum and maximum, or mid-point) converted into decimal degrees. Coordinate precision was likely variable among studies and is overall unknown. If studies did not report coordinates, we used the name given to the study site and location country to populate the coordinates using OpenCage (Salmon, 2018). OpenCage provides coordinates and a degree of confidence in the estimate, where 1 is low and 9 is high. For all coordinates were the confidence level fell below 7, we manually checked and if needed amended coordinates. When reported in the primary source, we also recorded the area (size) of the study site. For the study period in each record, we noted the start and end date of the population monitoring, and if available the corresponding population sizes at these dates. We captured the data collection and analysis methods from each source using several descriptors (Table S1). For studies that combined multiple methods, we precautionarily recorded the least robust approach. If we could not identify the method, the record was assigned ‘undefined’.

### Causes of change

Some sources tested or discussed the role of distinct factors to explain observed population changes. We recorded these factors reclassified into a modified version of the IUCN standardized classification schemes for Threats (v3.2) and Conservation Actions (v2.0), see Table S7. For each recorded factor we noted its effect (associated to increase or to decrease) and how this influence was determined. It is important to note that effects were not always negative for the threat scheme or positive for the conservation actions scheme. For example, urbanisation is listed under the threat scheme but has led to population increases in red fox *Vulpes vulpes* (Gloor *et al*., 2001). Finally, we note that factors not listed for a given record do not imply a threat or conservation action was not important or did not occur in that population, but simply that the factor was not mentioned in the primary literature.

### Validating records

Authors TJF and PC read the English and Spanish sources, respectively. TFJ entered all data. To validate the records and ensure quality control, 10% of the records were reviewed by an additional author (either PC or MGS). We selected the 10% sample with a random stratified approach to ensure each of the different formats of trends were reviewed e.g. percentage change, population time-series, and qualitative descriptions. TFJ then further scrutinised and double-checked records to detect errors in TFJs original work, that of the second readers (PC and MGS), and identify causes of discrepancies in data entry. We tested the reproducibility of our methods using the Grames & Elphick (2020) checklist and scored highly (Table S9).

## Results

From the 542 sources read in full, 232 did not contain the population trend information we required and were excluded from the dataset. Trends were excluded for a variety of reasons, examples include: the trend was simulated (N = 23), the trend referred to primary sources already captured in the dataset (N = 20), the trend described geographic distribution range change instead of abundance change (N = 6). Results from the validation step are reported in Supplementary: Validating records.

We identified and recorded 1122 population trends from the remaining 310 sources. These represented 50 (57%) of the studied species covering all four taxonomic families and 25 (69%) out of 36 genera (Figure 1). Some species had a single trend estimate, while we compiled 621 trend estimates for the top five species: gray wolf (*Canis lupus*), brown bear (*Ursus arctos*), grizzly bear (*Ursus americanus*), lion (*Panthera leo*) and Eurasian lynx (*Lynx lynx*). Many of the records represented populations within the northern hemisphere (Figure 2a), particularly in Europe (N = 384) and North America (N = 415), but there was also a cluster of records in East and Southern Africa (N = 170) – with records in 86 countries in total. We located very few records in Central, North and West Africa, Central and South America, or Northern Asia. The dataset includes records extending from 1726-2017 (Figure 2b), with the vast majority (92%) of trends starting after 1950.

**Figure 1.**
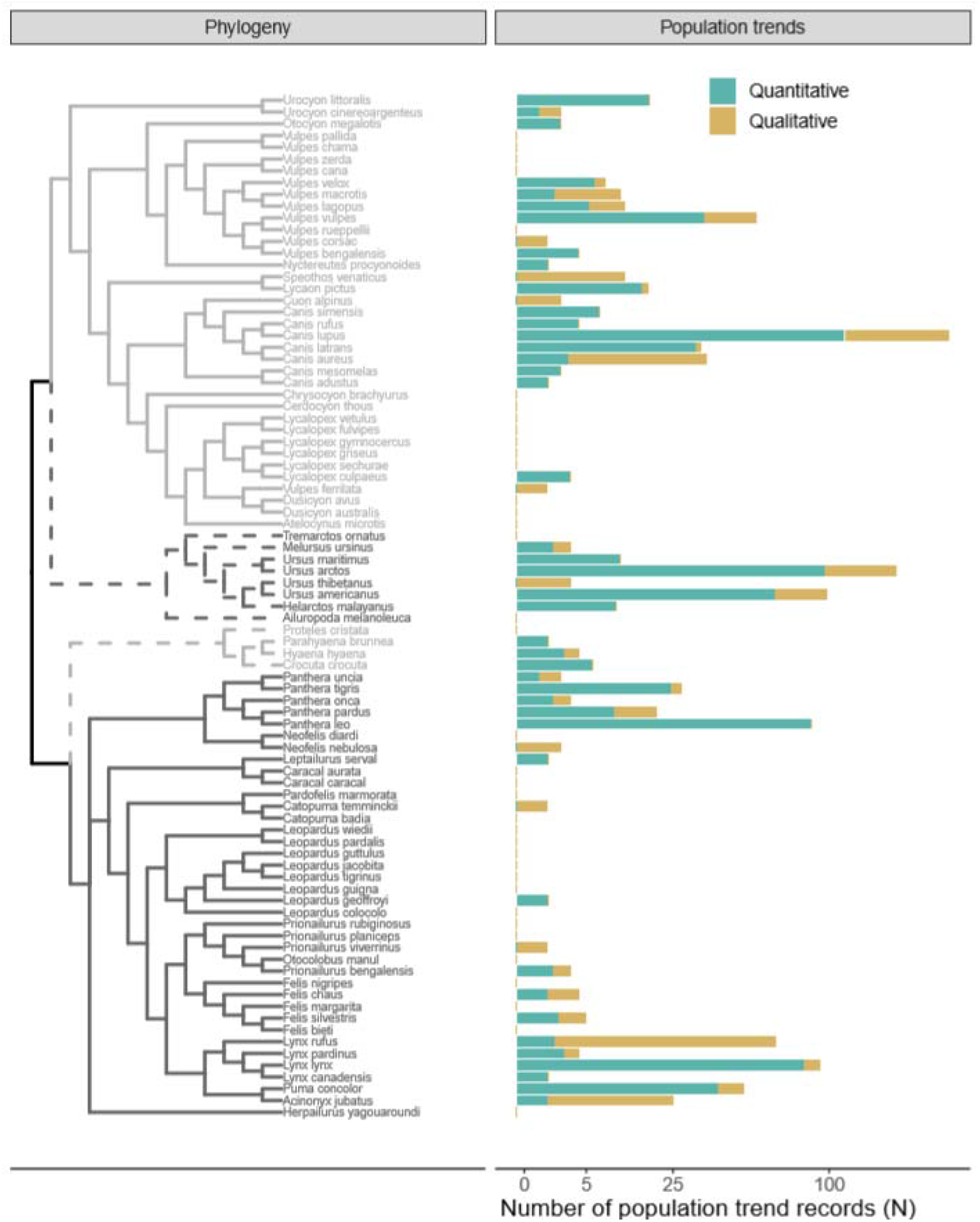
Number of population trend records per studied species, shown across the phylogeny. The tree represents four taxonomic families: Canidae (light grey – solid line), Ursidae (dark grey – dotted line), Hyaenidae (light grey – dotted line) and Felidae (dark grey – solid line). We show records for both quantitative (teal) and qualitative (gold) trends.

**Figure 2.**
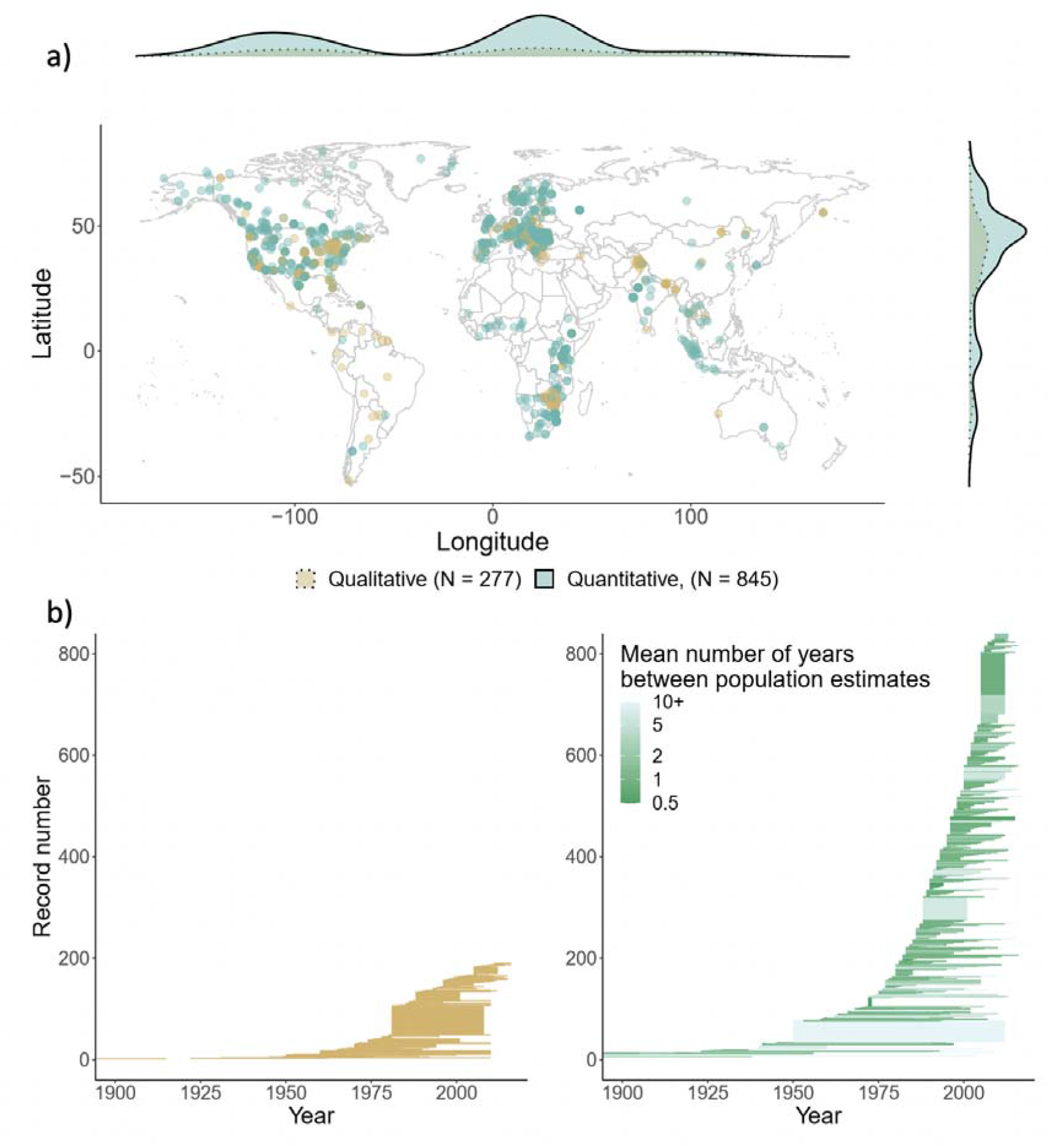
**a)** Location of study populations from which we compiled quantitative (teal) and qualitative (gold) trend records. Density plots indicate the frequency of the data points at varying latitudes and longitudes. Coordinates are decimal degrees. **b)** Distribution of qualitative (gold) and quantitative (green) population trend records between 1900-2017. Start and end date of each population trend record, ranked in ascending order of study start date. For the quantitative plot, we display the mean number of years between population estimates in each trend as a proxy for sampling effort, with darker green indicating greater sampling effort.

Most of the 1122 population trends represent quantitative estimates (N = 845), with a quarter (N = 277) providing only qualitative descriptors. The quantitative records collectively represent 6597 population size estimates. Most of the quantitative trends are recorded as a time-series of abundance values (63.9%), followed by population lambdas (17.4%), percentage change (7.5%), fold change (5.8%), and annual slope coefficients (5.4%).

## Discussion

We searched the literature to retrieve population trend records for 87 species of large carnivorans, and located 1122 estimates of population change representing 50 species. These records cover a wide temporal window (1726-2017) and represent diverse locations around the globe, although, there is temporal and spatial heterogeneity with more records in recent years and temperate areas of the Northern hemisphere. Our effort expands on and complements previous datasets for these species (as of September 2021, the Living Planet Index includes 465 trends across 39 species, and BioTIME includes 72 trends across 4 species) and thus, CaPTrends provides a valuable resource to address ecological questions, complete a more comprehensive assessment of population status for these species, and explore potential predictors of observed population changes (Johnson, *et al*., 2021)

Our dataset located additional time-series records not reported in the Living Planet Index, but also added less precise and qualitative descriptors which need to be interpreted with caution. For example, we found that studies that provided summarised quantitative metrics (e.g. annual population growth) did not always offer estimates of their error and thus, we could not extract uncertainty around the trend in all cases. This issue is even more emphasised in the qualitative descriptions (e.g. increase, decrease), where both the error and magnitude of the trend are unknown. However, if used cautiously, the lower resolution metrics could be important in addressing data gaps for species and locations for which high resolution population trend records are not available (WWF, 2016). This is particularly important, as these data gaps are most prevalent in biodiverse regions (WWF, 2016), which are experiencing the greatest negative-change in human footprint (Venter *et al*., 2016). Incorporating lower resolution metrics into models of biodiversity change could reduce some of these biases - providing a robust modelling approach is used. For example, in Johnson et al. (2021), trends are treated as a latent state, with qualitative estimates acting as an imperfect realisation of the trend.

## Usage notes

CaPTrends is presented as a relational dataset (Figure 3). The main file ‘captrends.csv’ includes all master data (e.g. unique id, species, location and time-frame), as well as all population data, except the population time-series. Time-series of population abundances and population changes are located in ‘ts_abundance.csv’ and ‘ts_change.csv’, respectively, both of which are linked to ‘captrends.csv’ through the ‘DataTableID’ field. ‘direction.csv’ also links to ‘captrends.csv’ through ‘DataTableID’ and describes positive and negative influences of each trend. Finally, ‘sources.csv’ links to ‘captrends.csv’ through ‘Citation_key’ and contains information on where the trend was sourced from (full reference). Comprehensive metadata is available for each of these datasets in the supplementary material.

**Figure 3.**
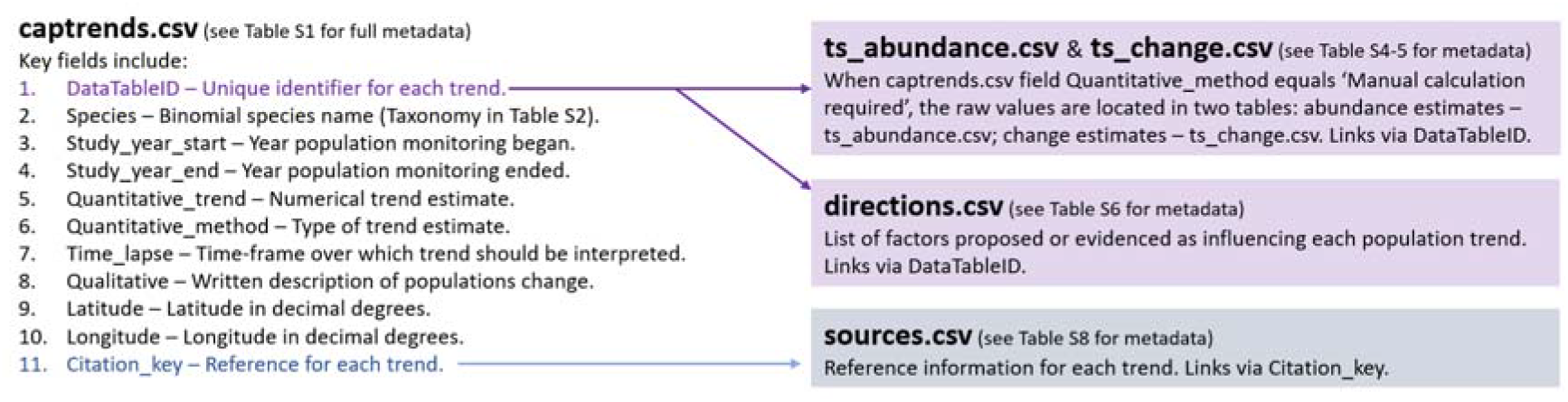
Diagram depicting relational database, including each datasets contents, and how each dataset is linked (arrows).

To support the use of this dataset, each population trend record has been annotated and labelled (Table S1). Much of this information would be helpful in filtering the dataset to exclude trends that are deemed of low quality or irrelevant to a given research question. For example, for investigating extinction risk, one may opt to remove data for invasive populations, which is just one of the indicator tags available for each trend.

This dataset may be analysed focusing on different descriptors. Including qualitative descriptors provides the most records but highest uncertainty. Focusing only on quantitative records reduces the scope and increase biases (not all species and areas are equally like to have quantitative records as shown in Figure 2). Approaches like data integration (Isaac *et al*., 2020), which can incorporate both data types, are likely to be least biased (spatially, temporally, and taxonomically).

## Acknowledgements

Thanks to Ella Coley, Jasmine Ashley, Jessica Marshall, Matthew Bemment, Monty Jefferson, Sarah Granger that assisted in data collection, and Julia Martínez Pardo that helped in project design. This work was funded by NERC (NE/P012345/1) and Royal Society (IE160539).

## Data accessibility

Data will be stored in an online repository. Location TBC

## Author contributions

All authors contributed to project design. TFJ entered the data with support from PC. Data was validated by MGS and PC. TFJ wrote the first draft of the manuscript, all authors contributed to revisions.

## References

Amano, T., Székely, T., Sandel, B., Nagy, S., Mundkur, T., Langendoen, T., Blanco, D., Soykan, C.U. & Sutherland, W.J. (2018) Successful conservation of global waterbird populations depends on effective governance. Nature.

Cardillo, M., Mace, G., Jones, K.E., Bininda-Emonds, O.R.P., Bielby, J., Sechrest, W., Orme, C.D.L. & Purvis, A. (2005) Multiple causes of high extinction risk in large mammal species. Science.

Chapron, G., Kaczensky, P., Linnell, J.D.C., von Arx, M., Huber, D., Andren, H., Lopez-Bao, J. V., Adamec, M., Alvares, F., Anders, O., Bal iauskas, L., Balys, V., Bed, P., Bego, F., Blanco, J.C., Breitenmoser, U., Broseth, H., Bufka, L., Bunikyte, R., Ciucci, P., Dutsov, A., Engleder, T., Fuxjager, C., Groff, C., Holmala, K., Hoxha, B., Iliopoulos, Y., Ionescu, O., Jeremi, J., Jerina, K., Kluth, G., Knauer, F., Kojola, I., Kos, I., Krofel, M., Kubala, J., Kunovac, S., Kusak, J., Kutal, M., Liberg, O., Maji, A., Mannil, P., Manz, R., Marboutin, E., Marucco, F., Melovski, D., Mersini, K., Mertzanis, Y., Mys ajek, R.W., Nowak, S., Odden, J., Ozolins, J., Palomero, G., Paunovi, M., Persson, J., Poto nik, H., Quenette, P.-Y., Rauer, G., Reinhardt, I., Rigg, R., Ryser, A., Salvatori, V., Skrbin ek, T., Stojanov, A., Swenson, J.E., Szemethy, L., Trajce, A., Tsingarska-Sedefcheva, E., Vaa, M., Veeroja, R., Wabakken, P., Wolfl, M., Wolfl, S., Zimmermann, F., Zlatanova, D. & Boitani, L. (2014) Recovery of large carnivores in Europe’s modern human-dominated landscapes. Science, 346, 1517–1519.

Dornelas, M., Antão, L.H., Moyes, F., Bates, A.E., Magurran, A.E., Adam, D., Akhmetzhanova, A.A., Appeltans, W., Arcos, J.M., Arnold, H., Ayyappan, N., Badihi, G., Baird, A.H., Barbosa, M., Barreto, T.E., Bässler, C., Bellgrove, A., Belmaker, J., Benedetti-Cecchi, L., Bett, B.J., Bjorkman, A.D., Błażewicz, M., Blowes, S.A., Bloch, C.P., Bonebrake, T.C., Boyd, S., Bradford, M., Brooks, A.J., Brown, J.H., Bruelheide, H., Budy, P., Carvalho, F., Castañeda-Moya, E., Chen, C.A., Chamblee, J.F., Chase, T.J., Siegwart Collier, L., Collinge, S.K., Condit, R., Cooper, E.J., Cornelissen, J.H.C., Cotano, U., Kyle Crow, S., Damasceno, G., Davies, C.H., Davis, R.A., Day, F.P., Degraer, S., Doherty, T.S., Dunn, T.E., Durigan, G., Duffy, J.E., Edelist, D., Edgar, G.J., Elahi, R., Elmendorf, S.C., Enemar, A., Ernest, S.K.M., Escribano, R., Estiarte, M., Evans, B.S., Fan, T.Y., Turini Farah, F., Loureiro Fernandes, L., Farneda, F.Z., Fidelis, A., Fitt, R., Fosaa, A.M., Daher Correa Franco, G.A., Frank, G.E., Fraser, W.R., García, H., Cazzolla Gatti, R., Givan, O., Gorgone-Barbosa, E., Gould, W.A., Gries, C., Grossman, G.D., Gutierréz, J.R., Hale, S., Harmon, M.E., Harte, J., Haskins, G., Henshaw, D.L., Hermanutz, L., Hidalgo, P., Higuchi, P., Hoey, A., Van Hoey, G., Hofgaard, A., Holeck, K., Hollister, R.D., Holmes, R., Hoogenboom, M., Hsieh C. hao, Hubbell, S.P., Huettmann, F., Huffard, C.L., Hurlbert, A.H., Macedo Ivanauskas, N., Janík, D., Jandt, U., Jażdżewska, A., Johannessen, T., Johnstone, J., Jones, J., Jones, F.A.M., Kang, J., Kartawijaya, T., Keeley, E.C., Kelt, D.A., Kinnear, R., Klanderud, K., Knutsen, H., Koenig, C.C., Kortz, A.R., Král, K., Kuhnz, L.A., Kuo, C.Y., Kushner, D.J., Laguionie-Marchais, C., Lancaster, L.T., Min Lee, C., Lefcheck, J.S., Lévesque, E., Lightfoot, D., Lloret, F., Lloyd, J.D., López-Baucells, A., Louzao, M., Madin, J.S., Magnússon, B., Malamud, S., Matthews, I., McFarland, K.P., McGill, B., McKnight, D., McLarney, W.O., Meador, J., Meserve, P.L., Metcalfe, D.J., Meyer, C.F.J., Michelsen, A., Milchakova, N., Moens, T., Moland, E., Moore, J., Mathias Moreira, C., Müller, J., Murphy, G., Myers-Smith, I.H., Myster, R.W., Naumov, A., Neat, F., Nelson, J.A., Paul Nelson, M., Newton, S.F., Norden, N., Oliver, J.C., Olsen, E.M., Onipchenko, V.G., Pabis, K., Pabst, R.J., Paquette, A., Pardede, S., Paterson, D.M., Pélissier, R., Peñuelas, J., Pérez-Matus, A., Pizarro, O., Pomati, F., Post, E., Prins, H.H.T., Priscu, J.C., Provoost, P., Prudic, K.L., Pulliainen, E., Ramesh, B.R., Mendivil Ramos, O., Rassweiler, A., Rebelo, J.E., Reed, D.C., Reich, P.B., Remillard, S.M., Richardson, A.J., Richardson, J.P., van Rijn, I., Rocha, R., Rivera-Monroy, V.H., Rixen, C., Robinson, K.P., Ribeiro Rodrigues, R., de Cerqueira Rossa-Feres, D., Rudstam, L., Ruhl, H., Ruz, C.S., Sampaio, E.M., Rybicki, N., Rypel, A., Sal, S., Salgado, B., Santos, F.A.M., Savassi-Coutinho, A.P., Scanga, S., Schmidt, J., Schooley, R., Setiawan, F., Shao, K.T., Shaver, G.R., Sherman, S., Sherry, T.W., Siciński, J., Sievers, C., da Silva, A.C., Rodrigues da Silva, F., Silveira, F.L., Slingsby, J., Smart, T., Snell, S.J., Soudzilovskaia, N.A., Souza, G.B.G., Maluf Souza, F., Castro Souza, V., Stallings, C.D., Stanforth, R., Stanley, E.H., Mauro Sterza, J., Stevens, M., Stuart-Smith, R., Rondon Suarez, Y., Supp, S., Yoshio Tamashiro, J., Tarigan, S., Thiede, G.P., Thorn, S., Tolvanen, A., Teresa Zugliani Toniato, M., Totland, Ø., Twilley, R.R., Vaitkus, G., Valdivia, N., Vallejo, M.I., Valone, T.J., Van Colen, C., Vanaverbeke, J., Venturoli, F., Verheye, H.M., Vianna, M., Vieira, R.P., Vrška, T., Quang Vu, C., Van Vu, L., Waide, R.B., Waldock, C., Watts, D., Webb, S., Wesołowski, T., White, E.P., Widdicombe, C.E., Wilgers, D., Williams, R., Williams, S.B., Williamson, M., Willig, M.R., Willis, T.J., Wipf, S., Woods, K.D., Woehler, E.J., Zawada, K. & Zettler, M.L. (2018) BioTIME: A database of biodiversity time series for the Anthropocene. Global Ecology and Biogeography, 27, 760–786.

Fritz, S.A., Bininda-Emonds, O.R.P. & Purvis, A. (2009) Geographical variation in predictors of mammalian extinction risk: Big is bad, but only in the tropics. Ecology Letters.

Gloor, S., Bontadina, F., Hegglin, D., Deplazes, P. & Breitenmoser, U. (2001) The rise of urban fox populations in Switzerland. Zeitschrift fur Saugetierkunde.

Gonzalez-Suarez, M., Gomez, A. & Revilla, E. (2013) Which intrinsic traits predict vulnerability to extinction depends on the actual threatening processes. Ecosphere, 4, 1–16.

González-Suárez, M. & Revilla, E. (2013) Variability in life-history and ecological traits is a buffer against extinction in mammals. Ecology Letters, 16, 242–251.

Grames, E.M. & Elphick, C.S. (2020) Use of study design principles would increase the reproducibility of reviews in conservation biology. Biological Conservation, 241, 108385.

IPBES (2019) Summary for policymakers of the global assessment report on biodiversity and ecosystem services of the Intergovernmental Science-Policy Platform on Biodiversity and Ecosystem Services,.

Isaac, N.J.B., Jarzyna, M.A., Keil, P., Dambly, L.I., Boersch-Supan, P.H., Browning, E., Freeman, S.N., Golding, N., Guillera-Arroita, G., Henrys, P.A., Jarvis, S., Lahoz-Monfort, J., Pagel, J., Pescott, O.L., Schmucki, R., Simmonds, E.G. & O’Hara, R.B. (2020) Data Integration for Large-Scale Models of Species Distributions. Trends in Ecology and Evolution.

Johnson, T.F., Isaac, N.J.B., Paviolo, A. & Gonzalez-Suarez, M. (2021) Socioeconomics drive population change in the world’s largest carnivore. bioRxiv - preprint.

Polaina, E., Revilla, E. & González-Suárez, M. (2016) Putting susceptibility on the map to improve conservation planning, an example with terrestrial mammals. Diversity and Distributions.

Ripple, W.J., Estes, J.A., Beschta, R.L., Wilmers, C.C., Ritchie, E.G., Hebblewhite, M., Berger, J., Elmhagen, B., Letnic, M., Nelson, M.P., Schmitz, O.J., Smith, D.W., Wallach, A.D. & Wirsing, A.J. (2014) Status and ecological effects of the world’s largest carnivores. Science, 343.

Rohatgi, A. (2015) WebPlotDigitizer.

Salmon, M. (2018) opencage: Interface to the OpenCage API.

Venter, O., Sanderson, E.W., Magrach, A., Allan, J.R., Beher, J., Jones, K.R., Possingham, H.P., Laurance, W.F., Wood, P., Fekete, B.M., Levy, M.A. & Watson, J.E.M. (2016) Sixteen years of change in the global terrestrial human footprint and implications for biodiversity conservation. Nature Communications.

Wilson, D.E. & Reeder, D.M. (2005) Mammal Species of the World,.

WWF (2016) Living Planet Report 2016: Risk and resilience in a new era, Gland, Switzerland.

WWF (2020) Living Planet Report 2020 - Bending the curve of biodiversity loss,.

