## Supplementary material for "CaPTrends: A global database of Carnivoran Population Trends"

**CaPTrends: A global database of population trends in large terrestrial Carnivorans – Supplementary material and dataset metadata**

Systematic search

Between September 2017 and January 2018, six individuals conducted a literature search for carnivore population trends in our four target families. The purpose of this search was to ensure there was sufficient data for a large data compilation effort, and to refine the fields we wished to collect. These semi-structured searches included two terms: a reference to the taxon (e.g. common name, scientific name, family, or order) and a population trend term (e.g. population trend*, declin*, increas*, recover*, conservation status, or population growth rate). These searches were conducted within Scopus, Web of Science and Google Scholar. The six individuals identified 80 peer-reviewed publications containing population trend data.

In February 2018, confident data were abundant enough for a large compilation effort, we conducted a structured search in Scopus and Web of Science looking for terms (see full search query in Supplementary material: Box S1) in the title, abstract and keywords discussing population trends in the world’s large terrestrial carnivorans. We considered the 87 species recognized in the four target families by the IUCN taxonomy (sourced February 2018). We queried the databases in English and Spanish. From this structured search, we found 3233 sources in English (reduced to 3060 after removing duplicates), and 30 sources in Spanish. Each of the Spanish sources were then read in full, but to further refine the English sources, each title was screened to remove sources not discussing the target taxa, e.g. references to ‘*tiger* shark’ were removed, this reduced the English sources to 1215.

In the remaining 1215 English sources, two readers read the same random sample of 50 abstracts and classified them stating whether a population trend for any of the 87 species is likely present or absent. Cohen’s kappa agreement between readers was substantial (68%), however it was clear some abstracts fell between the lines and could not be easily classified as containing a trend, or not. To account for this, one reader (TFJ) classified the remaining 1215 sources into five ordinal categories: 1 - explicitly mentions population trends of a target species (N = 539); 2 – trends of a target species are discussed but are not the primary focus of the manuscript (N = 155); 3 – trends are mentioned as part of the wider context (N = 164); 4 – population status is mentioned but the trend is not (N = 73); 5 – no population information mentioned (N = 284). Category 1 sources were then further refined to remove any sources discussing captive populations, simulated populations, or cases discussing trends in a non-target species (e.g. impact of lynx on hare population trends). This refining procedure left 468 category 1 sources, of which 32 had been identified in the unstructured search. After including the additional 48 sources identified in the unstructured search, we were left with 516 sources very likely to be discussing population trends in one of the target taxa. The entire refining process is summarised in Figure 1.

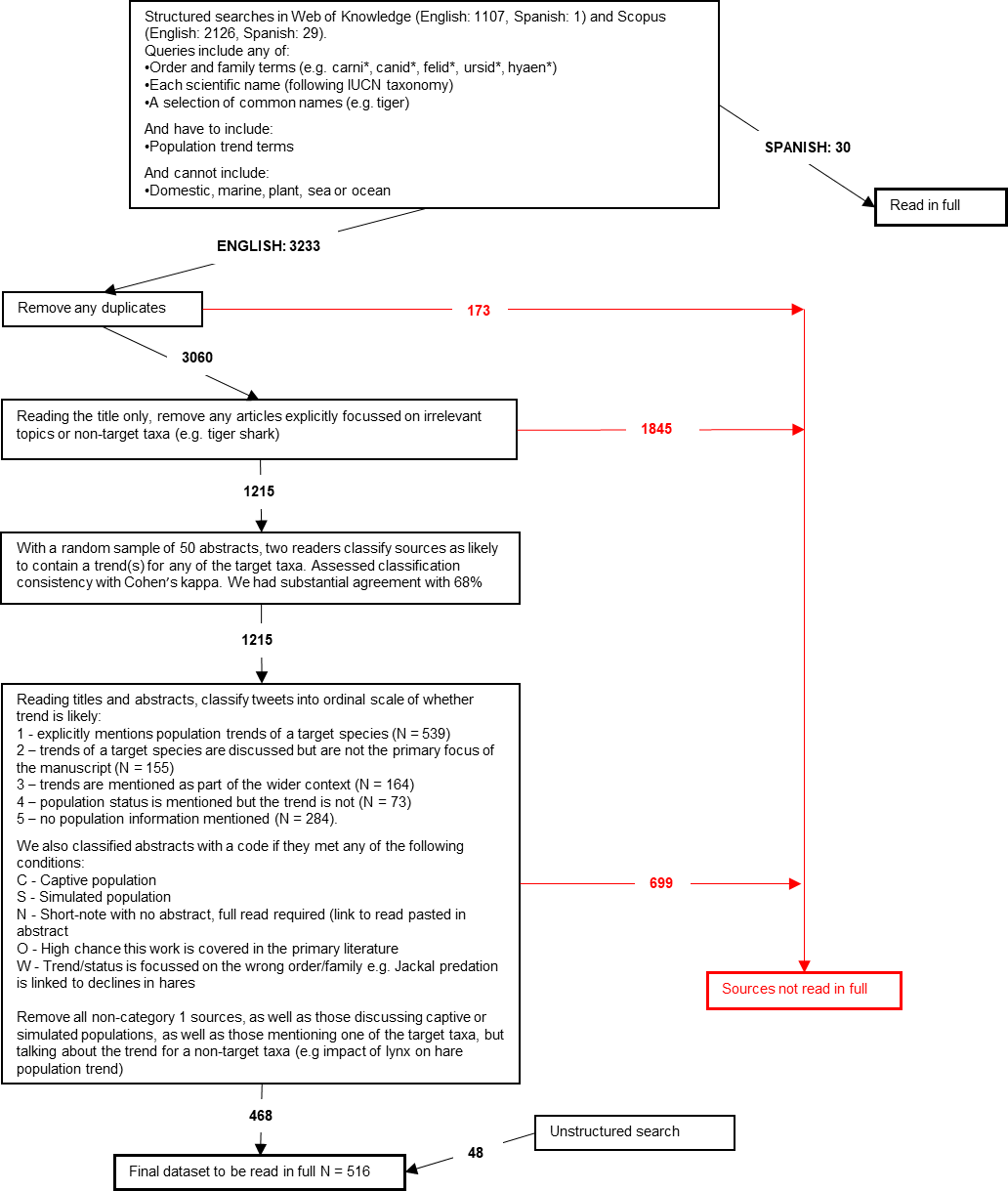

**Figure 1.** Data flow diagram specifying steps taken to identify publications which should be read in full. Black-arrows indicate sources taken forwards to next refining stage; the associated numbers indicate the count of publications brought through each step. The red-arrows indicate sources which at different steps were deemed irrelevant and not to be read in full.

Each of the highly relevant sources (N = 516) were read in full (no other categories were read in full). A selection of these category 1 sources was secondary literature providing population trend syntheses or compilations (van de Kerk *et al.*, 2013). In these cases, we located the original primary sources described in the secondary source and added them ad-hoc to category 1, increasing the overall number of sources to 536 which were to be read in full. Of these 536, we were unable to find the full text in ten cases, in five cases the primary data had already been captured, and in four cases the text was in language other than English or Spanish. After excluding these cases, we read the remaining 517 texts in full.

Validating records

Across the 10% (N = 31) of sources that underwent validation, TFJ and the second readers identified 46 population trends. The second readers located 40 and missed six. TFJ located 45 and missed one, which suggests that across all of the evaluated sources TFJ could have failed to detect ~2% of population trends. Further discrepancies were identified when TFJ re-scrutinised the 31 sources, and compared the original data entry to that of the second readers. The second readers misclassified more of the values than TFJ (7.4% vs 0%), produced more missing values (6.4% vs. 0.3%), and made more typos (0.5% vs. 0.3%). Despite these discrepancies, the results were qualitatively very similar in those trends identified by both TFJ and the second readers, with TFJ and the second readers producing the same trend value, same species, and similar locations e.g. TFJ and the second readers identified the same state or region in all cases. Furthermore, both TFJ and the second readers identified the same trends that should be treated cautiously and flagged with a warning in the dataset. All of this considered, the classification protocol was considered robust as TFJ, who entered the values in the full dataset, produced relatively few errors.

**Box S1.** Structured search queries in Web of Science and Scopus. Queries were developed and searched in both English and Spanish

**Web of Science (English):** TS=(( "carnivor*" OR "felid*" OR "canid*" OR "ursid*" OR "hyaen*" OR "Atelocynus microtis" OR "Canis adustus" OR "Canis aureus" OR "Canis latrans" OR "Canis lupus" OR "Canis mesomelas" OR "Canis rufus" OR "Canis simensis" OR "Cerdocyon thous" OR "Chrysocyon brachyurus" OR "Cuon alpinus" OR "Dusicyon australis" OR "Dusicyon avus" OR "Lycalopex culpaeus" OR "Lycalopex fulvipes" OR "Lycalopex griseus" OR "Lycalopex gymnocercus" OR "Lycalopex sechurae" OR "Lycalopex vetulus" OR "Lycaon pictus" OR "Nyctereutes procyonoides" OR "Otocyon megalotis" OR "Speothos venaticus" OR "Urocyon cinereoargenteus" OR "Urocyon littoralis" OR "Vulpes bengalensis" OR "Vulpes cana" OR "Vulpes chama" OR "Vulpes corsac" OR "Vulpes ferrilata" OR "Vulpes lagopus" OR "Vulpes macrotis" OR "Vulpes pallida" OR "Vulpes rueppellii" OR "Vulpes velox" OR "Vulpes vulpes" OR "Vulpes zerda" OR "Acinonyx jubatus" OR "Caracal aurata" OR "Caracal caracal" OR "Catopuma badia" OR "Catopuma temminckii" OR "Felis bieti" OR "Felis chaus" OR "Felis margarita" OR "Felis nigripes" OR "Felis silvestris" OR "Herpailurus yagouaroundi" OR "Leopardus colocolo" OR "Leopardus geoffroyi" OR "Leopardus guigna" OR "Leopardus guttulus" OR "Leopardus jacobita" OR "Leopardus pardalis" OR "Leopardus tigrinus" OR "Leopardus wiedii" OR "Leptailurus serval" OR "Lynx canadensis" OR "Lynx lynx" OR "Lynx pardinus" OR "Lynx rufus" OR "Neofelis diardi" OR "Neofelis nebulosa" OR "Otocolobus manul" OR "Panthera leo" OR "Panthera onca" OR "Panthera pardus" OR "Panthera tigris" OR "Panthera uncia" OR "Pardofelis marmorata" OR "Prionailurus bengalensis" OR "Prionailurus planiceps" OR "Prionailurus rubiginosus" OR "Prionailurus viverrinus" OR "Puma concolor" OR "Crocuta crocuta" OR "Hyaena hyaena" OR "Parahyaena brunnea" OR "Proteles cristata" OR "Ailuropoda melanoleuca" OR "Helarctos malayanus" OR "Melursus ursinus" OR "Tremarctos ornatus" OR "Ursus americanus" OR "Ursus arctos" OR "Ursus maritimus" OR "Ursus thibetanus" OR "Canis thous" OR "Canis brachyurus" OR "Canis alpinus" OR "Canis australis" OR "Pseudalopex culpaeus" OR "Pseudalopex fulvipes" OR "Pseudalopex griseus" OR "Pseudalopex gymnocercus" OR "Pseudalopex sechurae" OR "Pseudalopex vetulus" OR "Canis procyonoides" OR "Canis cinereoargenteus" OR "Vulpes littoralis" OR "Alopex lagopus" OR "Vulpes rueppelli " OR "Canis vulpes" OR "Fennecus zerda" OR "Felis jubata" OR "Profelis aurata" OR "Felis badia" OR "Pardofelis badia" OR "Felis temminckii" OR "Pardofelis temminckii" OR "Felis silvestris" OR "Felis yagouaroundi" OR "Herpailurus yagouaroundi" OR "Puma yagouaroundi" OR "Lynchailurus colocolo" OR "Oncifelis colocolo" OR "Oncifelis geoffroyi" OR "Oncifelis guigna" OR "Felis jacobita" OR "Oreailurus jacobita" OR "Oreailurus jacobitus" OR "Oreailurus jacobitus" OR "Caracal serval" OR "Felis nebulosa" OR "Felis manul" OR "Felis leo" OR "Felis onca" OR "Felis pardus" OR "Felis tigris" OR "Felis uncia" OR "Uncia uncia" OR "Felis concolor" OR "Hyaena brunnea" OR "Proteles cristatus" OR "Ursus melanoleucus" OR "Ursus malayanus" OR "Bradypus ursinus" OR "Ursus ornatus" OR "Thalarctos maritimus" OR "Aardwolf" OR "Fox" OR "Wolf" OR "Lynx" OR "Lion" OR "Leopard" OR "Caracal" OR "Bear" OR "Jackal" OR "Bobcat" OR "Puma" OR "Cougar" OR "Dhole" OR "Hyaena" OR "Cheetah" OR "Fennec" OR "Cat" OR "Dog" OR "Panda" OR "Margay" OR "Ocelot" OR "Tiger" ) AND ( "population trend*" OR "population dec*" OR "population increas*" OR "population recover*" OR "conservation status" OR "population growth" ) NOT ( "domestic" OR "marine" OR "plant" OR "sea" OR "ocean" ))

**Scopus (English):** TITLE-ABS-KEY ( "carnivor*" OR "felid*" OR "canid*" OR "ursid*" OR "hyaen*" OR "Atelocynus microtis" OR "Canis adustus" OR "Canis aureus" OR "Canis latrans" OR "Canis lupus" OR "Canis mesomelas" OR "Canis rufus" OR "Canis simensis" OR "Cerdocyon thous" OR "Chrysocyon brachyurus" OR "Cuon alpinus" OR "Dusicyon australis" OR "Dusicyon avus" OR "Lycalopex culpaeus" OR "Lycalopex fulvipes" OR "Lycalopex griseus" OR "Lycalopex gymnocercus" OR "Lycalopex sechurae" OR "Lycalopex vetulus" OR "Lycaon pictus" OR "Nyctereutes procyonoides" OR "Otocyon megalotis" OR "Speothos venaticus" OR "Urocyon cinereoargenteus" OR "Urocyon littoralis" OR "Vulpes bengalensis" OR "Vulpes cana" OR "Vulpes chama" OR "Vulpes corsac" OR "Vulpes ferrilata" OR "Vulpes lagopus" OR "Vulpes macrotis" OR "Vulpes pallida" OR "Vulpes rueppellii" OR "Vulpes velox" OR "Vulpes vulpes" OR "Vulpes zerda" OR "Acinonyx jubatus" OR "Caracal aurata" OR "Caracal caracal" OR "Catopuma badia" OR "Catopuma temminckii" OR "Felis bieti" OR "Felis chaus" OR "Felis margarita" OR "Felis nigripes" OR "Felis silvestris" OR "Herpailurus yagouaroundi" OR "Leopardus colocolo" OR "Leopardus geoffroyi" OR "Leopardus guigna" OR "Leopardus guttulus" OR "Leopardus jacobita" OR "Leopardus pardalis" OR "Leopardus tigrinus" OR "Leopardus wiedii" OR "Leptailurus serval" OR "Lynx canadensis" OR "Lynx lynx" OR "Lynx pardinus" OR "Lynx rufus" OR "Neofelis diardi" OR "Neofelis nebulosa" OR "Otocolobus manul" OR "Panthera leo" OR "Panthera onca" OR "Panthera pardus" OR "Panthera tigris" OR "Panthera uncia" OR "Pardofelis marmorata" OR "Prionailurus bengalensis" OR "Prionailurus planiceps" OR "Prionailurus rubiginosus" OR "Prionailurus viverrinus" OR "Puma concolor" OR "Crocuta crocuta" OR "Hyaena hyaena" OR "Parahyaena brunnea" OR "Proteles cristata" OR "Ailuropoda melanoleuca" OR "Helarctos malayanus" OR "Melursus ursinus" OR "Tremarctos ornatus" OR "Ursus americanus" OR "Ursus arctos" OR "Ursus maritimus" OR "Ursus thibetanus" OR "Canis thous" OR "Canis brachyurus" OR "Canis alpinus" OR "Canis australis" OR "Pseudalopex culpaeus" OR "Pseudalopex fulvipes" OR "Pseudalopex griseus" OR "Pseudalopex gymnocercus" OR "Pseudalopex sechurae" OR "Pseudalopex vetulus" OR "Canis procyonoides" OR "Canis cinereoargenteus" OR "Vulpes littoralis" OR "Alopex lagopus" OR "Vulpes rueppelli " OR "Canis vulpes" OR "Fennecus zerda" OR "Felis jubata" OR "Profelis aurata" OR "Felis badia" OR "Pardofelis badia" OR "Felis temminckii" OR "Pardofelis temminckii" OR "Felis silvestris" OR "Felis yagouaroundi" OR "Herpailurus yagouaroundi" OR "Puma yagouaroundi" OR "Lynchailurus colocolo" OR "Oncifelis colocolo" OR "Oncifelis geoffroyi" OR "Oncifelis guigna" OR "Felis jacobita" OR "Oreailurus jacobita" OR "Oreailurus jacobitus" OR "Oreailurus jacobitus" OR "Caracal serval" OR "Felis nebulosa" OR "Felis manul" OR "Felis leo" OR "Felis onca" OR "Felis pardus" OR "Felis tigris" OR "Felis uncia" OR "Uncia uncia" OR "Felis concolor" OR "Hyaena brunnea" OR "Proteles cristatus" OR "Ursus melanoleucus" OR "Ursus malayanus" OR "Bradypus ursinus" OR "Ursus ornatus" OR "Thalarctos maritimus" OR "Aardwolf" OR "Fox" OR "Wolf" OR "Lynx" OR "Lion" OR "Leopard" OR "Caracal" OR "Bear" OR "Jackal" OR "Bobcat" OR "Puma" OR "Cougar" OR "Dhole" OR "Hyaena" OR "Cheetah" OR "Fennec" OR "Cat" OR "Dog" OR "Panda" OR "Margay" OR "Ocelot" OR "Tiger" ) AND ( "population trend*" OR "population dec*" OR "population increas*" OR "population recover*" OR "conservation status" OR "population growth" ) AND NOT ( "domestic" OR "marine" OR "plant" OR "sea" OR "ocean" )

**Web of Science (Spanish):** TS = (( "carnivor*" OR "felid*" OR "canid*" OR "ursid*" OR "hyaen*" OR "Atelocynus microtis" OR "Canis adustus" OR "Canis aureus" OR "Canis latrans" OR "Canis lupus" OR "Canis mesomelas" OR "Canis rufus" OR "Canis simensis" OR "Cerdocyon thous" OR "Chrysocyon brachyurus" OR "Cuon alpinus" OR "Dusicyon australis" OR "Dusicyon avus" OR "Lycalopex culpaeus" OR "Lycalopex fulvipes" OR "Lycalopex griseus" OR "Lycalopex gymnocercus" OR "Lycalopex sechurae" OR "Lycalopex vetulus" OR "Lycaon pictus" OR "Nyctereutes procyonoides" OR "Otocyon megalotis" OR "Speothos venaticus" OR "Urocyon cinereoargenteus" OR "Urocyon littoralis" OR "Vulpes bengalensis" OR "Vulpes cana" OR "Vulpes chama" OR "Vulpes corsac" OR "Vulpes ferrilata" OR "Vulpes lagopus" OR "Vulpes macrotis" OR "Vulpes pallida" OR "Vulpes rueppellii" OR "Vulpes velox" OR "Vulpes vulpes" OR "Vulpes zerda" OR "Acinonyx jubatus" OR "Caracal aurata" OR "Caracal caracal" OR "Catopuma badia" OR "Catopuma temminckii" OR "Felis bieti" OR "Felis chaus" OR "Felis margarita" OR "Felis nigripes" OR "Felis silvestris" OR "Herpailurus yagouaroundi" OR "Leopardus colocolo" OR "Leopardus geoffroyi" OR "Leopardus guigna" OR "Leopardus guttulus" OR "Leopardus jacobita" OR "Leopardus pardalis" OR "Leopardus tigrinus" OR "Leopardus wiedii" OR "Leptailurus serval" OR "Lynx canadensis" OR "Lynx lynx" OR "Lynx pardinus" OR "Lynx rufus" OR "Neofelis diardi" OR "Neofelis nebulosa" OR "Otocolobus manul" OR "Panthera leo" OR "Panthera onca" OR "Panthera pardus" OR "Panthera tigris" OR "Panthera uncia" OR "Pardofelis marmorata" OR "Prionailurus bengalensis" OR "Prionailurus planiceps" OR "Prionailurus rubiginosus" OR "Prionailurus viverrinus" OR "Puma concolor" OR "Crocuta crocuta" OR "Hyaena hyaena" OR "Parahyaena brunnea" OR "Proteles cristata" OR "Ailuropoda melanoleuca" OR "Helarctos malayanus" OR "Melursus ursinus" OR "Tremarctos ornatus" OR "Ursus americanus" OR "Ursus arctos" OR "Ursus maritimus" OR "Ursus thibetanus" OR "Canis thous" OR "Canis brachyurus" OR "Canis alpinus" OR "Canis australis" OR "Pseudalopex culpaeus" OR "Pseudalopex fulvipes" OR "Pseudalopex griseus" OR "Pseudalopex gymnocercus" OR "Pseudalopex sechurae" OR "Pseudalopex vetulus" OR "Canis procyonoides" OR "Canis cinereoargenteus" OR "Vulpes littoralis" OR "Alopex lagopus" OR "Vulpes rueppelli " OR "Canis vulpes" OR "Fennecus zerda" OR "Felis jubata" OR "Profelis aurata" OR "Felis badia" OR "Pardofelis badia" OR "Felis temminckii" OR "Pardofelis temminckii" OR "Felis silvestris" OR "Felis yagouaroundi" OR "Herpailurus yagouaroundi" OR "Puma yagouaroundi" OR "Lynchailurus colocolo" OR "Oncifelis colocolo" OR "Oncifelis geoffroyi" OR "Oncifelis guigna" OR "Felis jacobita" OR "Oreailurus jacobita" OR "Oreailurus jacobitus" OR "Oreailurus jacobitus" OR "Caracal serval" OR "Felis nebulosa" OR "Felis manul" OR "Felis leo" OR "Felis onca" OR "Felis pardus" OR "Felis tigris" OR "Felis uncia" OR "Uncia uncia" OR "Felis concolor" OR "Hyaena brunnea" OR "Proteles cristatus" OR "Ursus melanoleucus" OR "Ursus malayanus" OR "Bradypus ursinus" OR "Ursus ornatus" OR "Thalarctos maritimus" OR "Borochi" OR "Chacal" OR "Zorro" OR "Gato" OR "León" OR "Manigordo" OR "Manul" OR "Mitzli" OR "Ocelote" OR "Onza" OR "Oso" OR "Pacha" OR "Panda" OR "Pantera" OR "Perro" OR "Renard" OR "Tigre" OR "Tigrillo" OR "Tirica" OR "Umba" OR "Yaguar*" OR "Jaguar" OR "Lince" OR "Lobo" OR "Aguará" ) AND ( "tendencia poblacional" OR "declinación poblacional" OR "incremento poblacional" OR "recuperación poblacional" OR "estado de conservación" OR "tasa de incremento poblacional" ) NOT ( "Doméstic*" OR "Marin*" OR "Planta" OR "Océano" ))

**Scopus (Spanish):** TITLE-ABS-KEY ( "carnivor*" OR "felid*" OR "canid*" OR "ursid*" OR "hyaen*" OR "Atelocynus microtis" OR "Canis adustus" OR "Canis aureus" OR "Canis latrans" OR "Canis lupus" OR "Canis mesomelas" OR "Canis rufus" OR "Canis simensis" OR "Cerdocyon thous" OR "Chrysocyon brachyurus" OR "Cuon alpinus" OR "Dusicyon australis" OR "Dusicyon avus" OR "Lycalopex culpaeus" OR "Lycalopex fulvipes" OR "Lycalopex griseus" OR "Lycalopex gymnocercus" OR "Lycalopex sechurae" OR "Lycalopex vetulus" OR "Lycaon pictus" OR "Nyctereutes procyonoides" OR "Otocyon megalotis" OR "Speothos venaticus" OR "Urocyon cinereoargenteus" OR "Urocyon littoralis" OR "Vulpes bengalensis" OR "Vulpes cana" OR "Vulpes chama" OR "Vulpes corsac" OR "Vulpes ferrilata" OR "Vulpes lagopus" OR "Vulpes macrotis" OR "Vulpes pallida" OR "Vulpes rueppellii" OR "Vulpes velox" OR "Vulpes vulpes" OR "Vulpes zerda" OR "Acinonyx jubatus" OR "Caracal aurata" OR "Caracal caracal" OR "Catopuma badia" OR "Catopuma temminckii" OR "Felis bieti" OR "Felis chaus" OR "Felis margarita" OR "Felis nigripes" OR "Felis silvestris" OR "Herpailurus yagouaroundi" OR "Leopardus colocolo" OR "Leopardus geoffroyi" OR "Leopardus guigna" OR "Leopardus guttulus" OR "Leopardus jacobita" OR "Leopardus pardalis" OR "Leopardus tigrinus" OR "Leopardus wiedii" OR "Leptailurus serval" OR "Lynx canadensis" OR "Lynx lynx" OR "Lynx pardinus" OR "Lynx rufus" OR "Neofelis diardi" OR "Neofelis nebulosa" OR "Otocolobus manul" OR "Panthera leo" OR "Panthera onca" OR "Panthera pardus" OR "Panthera tigris" OR "Panthera uncia" OR "Pardofelis marmorata" OR "Prionailurus bengalensis" OR "Prionailurus planiceps" OR "Prionailurus rubiginosus" OR "Prionailurus viverrinus" OR "Puma concolor" OR "Crocuta crocuta" OR "Hyaena hyaena" OR "Parahyaena brunnea" OR "Proteles cristata" OR "Ailuropoda melanoleuca" OR "Helarctos malayanus" OR "Melursus ursinus" OR "Tremarctos ornatus" OR "Ursus americanus" OR "Ursus arctos" OR "Ursus maritimus" OR "Ursus thibetanus" OR "Canis thous" OR "Canis brachyurus" OR "Canis alpinus" OR "Canis australis" OR "Pseudalopex culpaeus" OR "Pseudalopex fulvipes" OR "Pseudalopex griseus" OR "Pseudalopex gymnocercus" OR "Pseudalopex sechurae" OR "Pseudalopex vetulus" OR "Canis procyonoides" OR "Canis cinereoargenteus" OR "Vulpes littoralis" OR "Alopex lagopus" OR "Vulpes rueppelli " OR "Canis vulpes" OR "Fennecus zerda" OR "Felis jubata" OR "Profelis aurata" OR "Felis badia" OR "Pardofelis badia" OR "Felis temminckii" OR "Pardofelis temminckii" OR "Felis silvestris" OR "Felis yagouaroundi" OR "Herpailurus yagouaroundi" OR "Puma yagouaroundi" OR "Lynchailurus colocolo" OR "Oncifelis colocolo" OR "Oncifelis geoffroyi" OR "Oncifelis guigna" OR "Felis jacobita" OR "Oreailurus jacobita" OR "Oreailurus jacobitus" OR "Oreailurus jacobitus" OR "Caracal serval" OR "Felis nebulosa" OR "Felis manul" OR "Felis leo" OR "Felis onca" OR "Felis pardus" OR "Felis tigris" OR "Felis uncia" OR "Uncia uncia" OR "Felis concolor" OR "Hyaena brunnea" OR "Proteles cristatus" OR "Ursus melanoleucus" OR "Ursus malayanus" OR "Bradypus ursinus" OR "Ursus ornatus" OR "Thalarctos maritimus" OR "Borochi" OR "Chacal" OR "Zorro" OR "Gato" OR "León" OR "Manigordo" OR "Manul" OR "Mitzli" OR "Ocelote" OR "Onza" OR "Oso" OR "Pacha" OR "Panda" OR "Pantera" OR "Perro" OR "Renard" OR "Tigre" OR "Tigrillo" OR "Tirica" OR "Umba" OR "Yaguar*" OR "Jaguar" OR "Lince" OR "Lobo" OR "Aguará" ) AND ( "tendencia poblacional" OR "declinación poblacional" OR "incremento poblacional" OR "recuperación poblacional" OR "estado de conservación" OR "tasa de incremento poblacional" ) AND NOT ( "Doméstic*" OR "Marin*" OR "Planta" OR "Océano" ) AND ( LIMIT-TO ( LANGUAGE , "Spanish" ))

**Table S1.** Description of fields in the table captrends.csv. This is the core dataset with all master information and is the foundation that links to the other relational databases: timeseries.csv, direction.csv, and sources.csv. ‘Data type’ describes the format of the data, for categorical fields the selection options are underlined with its description italicised.

| **Field** | **Description** | **Data type** |
| --- | --- | --- |
| DataTableID | Unique numerical code for each population trend record. Matches with tables: ts_abundance.csv, ts_change.csv, direction.csv. | Character |
| Species | Binomial species name following IUCN taxonomy | Character [populated from Table S2] |
| Sub_species | Subspecies as listed within the source | Character |
| Citation_key | Unique alphanumerical code for each source to match with table sources.csv | Character |
| Spatial_locality | If papers have trends split into different sites, each site is given its own spatial unique numerical code | Numeric |
| Temporal_locality | If papers have trends split into different time points (e.g. 1980 - 1990, and 1990 - 2000), each consecutive time series is given its own temporal numerical code | Numeric |
| Locality_name | Name of study site as described in the primary source | Character |
| Singular_country | Country where studied population occurs following ISO3166 naming standards as of 2018 (e.g. source mentions Soviet Union and coordinates indicate Russia, Russia was recorded). | Character [populated from Table S3] |
| Multiple_countries | When studied population overlaps multiple countries, each country is included in a list separated with semi-colons. Country names follow ISO3166 standards. | Character [populated from supplementary: Country list] |
| Wider_population | Further information about the study site e.g. name of the region, state or national park. | Character |
| Locality_area | Numeric estimate of the study site area | Numeric |
| Locality_area_units | Units in which area of study site ‘Locality_area’ is reported. Categories:  *Hectares: Area where the population was studied (recorded in hectares)*  *Km^2^: Area where the population was studied (recorded in square kilometres)* | Categorical |
| Study_year_start | Year of first population size estimate | Numeric |
| Study_year_end | Year of final population size estimate | Numeric |
| N_observations | Number of population size estimates used to derive the trend. For quantitative population trends, the minimum value is 2. For records which include the complete time-series the value is missing but can be extracted from the ts_abundance.csv and ts_change.csv tables. For qualitative trends this value regularly equals one or zero, as there are many cases in the qualitative trends where only one or zero population estimates are made, and the assessment of the trend is more subjective. For matrix models, this value represents the number of sampling years, rather than the number of population size estimates. | Numeric |
| Field_method | Field method for deriving population size estimates or demographic information. Categories:  *Individuals identified: All individuals of a population were identified.*  *Systematic – direct: Monitoring approach is systematic (not-opportunistic), is not clearly prone to spatial or temporal bias, and involves direct observations of the animal (either alive or dead) e.g. through camera-trap grids or road-transects.*  *Systematic – indirect: Monitoring approach is systematic (not-opportunistic), is not clearly prone to spatial or temporal bias, and involves indirect observations of the animal e.g. footprint, audio calls, fur traps.*  *Systematic -undefined: Monitoring approach is systematic (not-opportunistic) and is not clearly prone to spatial or temporal bias but the actual method of making observations is unclear or a mix of direct and indirect.*  *Unsystematic – direct: Monitoring approach is opportunistic or not completely systematic and is at least partially prone to spatial or temporal bias; also involves direct observations of the animal (either alive or dead) e.g. through camera-trap grids or road-transects.*  *Unsystematic – indirect: Monitoring approach is opportunistic or not completely systematic and is at least partially prone to spatial or temporal bias; also involves indirect observations of the animal e.g. footprint, audio calls, fur traps.*  *Unsystematic - undefined: Monitoring approach is opportunistic or not completely systematic and is at least partially prone to spatial or temporal bias; also the actual method of making observations is unclear or a mix of direct and indirect.*  *Undefined: Population monitoring method poorly defined or does not meet one of the above criteria*. | Categorical |
| Modelling_method | Analysis method for deriving population estimates or demographic information. Categories:  *Model derived abundance/density: Statistical model used to convert field data into population abundance or density estimates.*  *Model occupancy: Statistical model used to convert occupancy field data into population abundance or density estimates.*  *Matrix modelling: Statistical model to estimate population change using demographic parameters.*  *Total count: Total population size is known, no need for statistical inference of abundance.*  *Relative abundance: Statistical approach to control for different sampling effort in detection events e.g. relative abundance.*  *Field values: Raw field data presented, no statistical modelling used to control for differences in sampling effort, observers etc.*  *Undefined: Approach for estimating population size is unclear or not explained, or does not clearly fall into any other category.* | Categorical |
| Population_metric | Type of population size measurement. Categories:  *Abundance: Estimates of the number of individuals in the population.*  *Density: Estimate of the number of individuals per unit of area. Units defined by variable 'Density_scale'.*  *Other: Estimate of the population size in alternate units e.g. relative abundance.* | Categorical |
| Density_scale | Units of population_metric when reported as *Density*. | Character |
| Population_start | Population size estimate in the first recorded year (as listed in field ‘Study_year_start’). Type of estimate described in field ‘Population_metric ‘ | Numeric |
| PS_dispersion_estimate | Estimate of dispersion or uncertainty in the population size value provided in field ‘Population_start’. Values entered here when they are provided as single estimate (e.g., SE or SD) Type of estimate described in field ‘PS_PE_dispersion_description’ | Numeric |
| PS_dispersion_lower | Estimate of dispersion or uncertainty in the population size value provided in field ‘Population_start’. Values entered here when they are provided as a lower bounded estimate (e.g., range or confidence intervals) Type of estimate described in field ‘PS_PE_dispersion_description’ | Numeric |
| PS_dispersion_upper | Estimate of dispersion or uncertainty in the population size value provided in field ‘Population_start’. Values entered here when they are provided as an upper bounded estimate (e.g., range or confidence intervals) Type of estimate described in field ‘PS_PE_dispersion_description’ | Numeric |
| Population_end | Population size estimate in the last recorded year (as listed in field ‘Study_year_end’). Type of estimate described in field ‘Population_metric ‘ | Numeric |
| PE_dispersion_estimate | Estimate of dispersion or uncertainty in the population size value provided in field ‘Population_end’. Values entered here when they are provided as single estimate (e.g., SE or SD) Type of estimate described in field ‘PS_PE_dispersion_description’ | Numeric |
| PE_dispersion_lower | Estimate of dispersion or uncertainty in the population size value provided in field ‘Population_end’. Values entered here when they are provided as a lower bounded estimate (e.g., range or confidence intervals) Type of estimate described in field ‘PS_PE_dispersion_description’ | Numeric |
| PE_dispersion_upper | Estimate of dispersion or uncertainty in the population size value provided in field ‘Population_end’. Values entered here when they are provided as an upper bounded estimate (e.g., range or confidence intervals) Type of estimate described in field ‘PS_PE_dispersion_description’ | Numeric |
| PS_PE_dispersion_description | Type of dispersion or uncertainty estimate(s) in population size values. Categories:  *SD: Standard deviation.*  *SE: Standard error.*  *Range: Minimum and maximum estimates.*  *90% CI: 90% confidence intervals.*  *95% CI: 95% confidence intervals.*  *Bayesian 90% CI: 90% credible intervals derived through Bayesian sampling.* | Categorical |
| Quantitative_trend | Numerical estimate of change in population size. Type of estimate described in field ‘Quantitative_method’. | Numeric |
| Quantitative_method | Type of population trend metric provided in field ‘Quantitative_trend’. Categories:  *Manual calculation required: complete time series available in the table [timeseries.csv]. Data fall into two categories: 1) estimates of abundance at different time points. 2) Estimates of change in abundance (e.g. population lambda, or percent change) at different time points. See metadata: timeseries.csv for more detail.*  *Lambda: finite rate of population change (lambda=1 represents a stable trend). Lambdas were estimated using different methods including ratio of abundance between two time intervals (N_t+1_/N_t_), different demographic models, or as the exponential of an R-trend coefficient.*  *R-trend:* *instantaneous rate of population change. Values were calculated with different methods but most frequently using a log-regression model of population size (R-trend = 0 represents a stable trend).*  *Percentage change: change in population size between two time points (100 is stable) [formula = (N_t+1_/N_t_) * 100].*  *Fold change: change in population size between two time points (1 is stable) [formula = (N_t+1_/N_t_)].*  *Qualitative only: only a verbal description of population change was available*. | Categorical |
| Other_quantitative_descriptor | Additional notes and comments about the quantitative descriptor extracted during data compilation to explain less-clear cases. | Character |
| Dispersion_description | Type of estimate of dispersion or uncertainty provided for the population trend metric. Estimate of dispersion provided in field ‘Dispersion_estimate’. Categories:  *VAR: Variance.*  *SD: Standard deviation.*  *SE: Standard error.*  *Range: Minimum and maximum estimates.*  *90% CI: 90% confidence intervals.*  *95% CI: 95% confidence intervals.*  *Bayesian 90% CI: 90% credible intervals derived through Bayesian sampling*. | Categorical |
| Dispersion_estimate | Estimate of dispersion or uncertainty for population trend (provided in field ‘Quantitative_trend field’). Type of uncertainty/dispersion described in field ‘PS_PE_dispersion_description’ | Numeric |
| Dispersion_lower | Estimate of lower bound dispersion or uncertainty (e.g., confidence intervals or range) for population trend (provided in field ‘Quantitative_trend’). Type of uncertainty/dispersion described in field ‘Dispersion_description’ | Numeric |
| Dispersion_upper | Estimate of upper bound dispersion or uncertainty (e.g., confidence intervals or range) for population trend (provided in field ‘Quantitative_trend’). Type of uncertainty/dispersion described in field ‘Dispersion_description’ | Numeric |
| Significance_reported | Descriptor of whether statistical significance in population trend was tested. Categories:  *NA: not reported or not relevant.*  *Yes: test statistic and/or significance level reported.* | Categorical |
| Test_statistic | Value of the statistic (e.g. z, t, or F value) used to describe significance in population trend when available. | Numeric |
| Significance | P-value associated to the ‘Test_statistic’ used to describe significance in population trend when available. | Numeric |
| Significant_trend | Binary descriptor of whether, if statistically tested, the population trend was found to be significantly increasing or declining. Categories:  *TRUE: trend was significant.*  *FALSE: trend was not-significant* | Categorical |
| Time_lapse | Timeframe (in years) at which Quantitative_trend should be interpreted e.g. a 10-year study may describe the annual finite rate of change (lambda), as its annual the Time_lapse would equal 1. However, some lambda’s are measured at 0.5 year or 10 year scale, so the metric is used to scale the Quanittative_trend to a standard time-frame. This value equals NA when the Quantitative_method is Qualitative only or a Manual trend estimate. | Numeric |
| Qualitative | Verbal description of population change as provided by the primary sources/publications. Categories:  *Increase: trend described as increasing, or recovering, or something synonymous.*  *Stable: trend described as stable or exhibiting no population change, or something synonymous.*  *Decrease: trend as described decreasing, declining, or reducing, or something synonymous.*  *Varied: trend described as showing both increases and decreases at different time periods, but crucially, the first and the last population estimates are similar.* | Category |
| Other_driver_of_trend | Factors described in source as influencing population trends but which could not be captured by threat or conservation actions schema | Character |
| Comment | Additional notes and comments extracted during data compilation. | Character |
| Possible_issues | Description of issues that may limit use or interpretation of the trend e.g. author may describe the trend estimate as inaccurate. | Character |
| Genetic_data | Binary descriptor of whether the population trend was derived from genetic information. Categories:  *1: yes*  *NA: no* | Numeric-binary |
| Harvest_data | Binary descriptor of whether the population trend was derived from harvest information e.g. number of individuals hunted. Categories:  *1: yes*  *NA: no* | Numeric-binary |
| Invasive_species | Binary descriptor of whether the studied population was non-native to the study site. Categories:  *1: yes*  *NA: no* | Numeric-binary |
| Record_labelled_inaccurate | Binary descriptor of whether the population trend was described as inaccurate in the source. Categories:  *1: yes*  *NA: no* | Numeric-binary |
| Asymptotic_growth | Binary descriptor of whether the population trend described asymptotic or observed growth. Categories:  *1: yes*  *NA: no* | Numeric-binary |
| Metric_unusual | Binary descriptor of whether the population trend was reported in an unconventional way. Categories:  *1: yes*  *NA: no* | Numeric-binary |
| Peer_review | Binary descriptor of whether the source has been published after peer-reviewed. Categories:  *1: no*  *NA: yes* | Numeric-binary |
| Date_missing | Binary descriptor of whether any of the date values are missing (Study_year_start or Study_year_end). Categories:  *1: yes*  *NA: no* | Numeric-binary |
| Latitude | Latitudinal centroid in decimal degrees of the study site/population | Numeric |
| Longitude | Longitudinal centroid in decimal degrees of the study site/population | Numeric |
| Source | Source of the coordinates. Categories:  *Georeferenced – automatically: obtained from OpenCage georeferencer using locality name and country from the source.*  *Georeferenced - manually adjusted: obtained from OpenCage georeferencer using locality name and country, but coordinates were inaccurate so were manually corrected.*  *Within study - calculated centroid: Coordinates included in the source as extent ranges from which the centroid was calculated.*  *Within study - reported centroid: centroid reported in the source.* | Categorical |
| Coordinate_comment | Process for reviewing coordinates that were georeferenced. Categories:  *Checked - location is approximate: georeferenced coordinates were checked and the precise location could not be found. Coordinates approximated manually.*  *Checked - Location refined: georeferenced coordinates were checked and the deemed inaccurate, so were manually adjusted.*  *Checked - Original is robust: georeferenced coordinates were checked and deemed robust.*  *Not checked - Record appears robust: georeferenced coordinates had high a confidence value (greater than or equal to 7) and so were not checked.*  *NA – coordinates not checked as they were extracted from the primary source.* | Categorical |

**Table S2.** Reference table for captrends.csv ‘Species’ field. Includes binomial species names for four target families (Canidae, Felidae, Hyaenidae, and Ursidea) within the order Carnivora. These species names follow the IUCN species list/taxonomy (downloaded in 2018), but we also provide comparison to the common mammalian reference taxonomy of (Wilson & Reeder, 2005). The comment column describes any discrepancies in these taxonomies to facilitate future dataset use.

| **Family** | **Species (IUCN)** | **Species (WR2005)** | **Comment** |
| --- | --- | --- | --- |
| *CANIDAE* | *Atelocynus microtis* | *Atelocynus microtis* |  |
|  | *Canis adustus* | *Canis adustus* |  |
|  | *Canis aureus* | *Canis aureus* |  |
|  | *Canis latrans* | *Canis latrans* |  |
|  | *Canis lupus* | *Canis lupus* |  |
|  | *Canis mesomelas* | *Canis mesomelas* |  |
|  | *Canis rufus* | *NA* | Considered a sub-species of *Canis lupus* in WR2005 |
|  | *Canis simensis* | *Canis simensis* |  |
|  | *Cerdocyon thous* | *Cerdocyon thous* |  |
|  | *Chrysocyon brachyurus* | *Chrysocyon brachyurus* |  |
|  | *Cuon alpinus* | *Cuon alpinus* |  |
|  | *Dusicyon australis* | *Dusicyon australis* |  |
|  | *Dusicyon avus* | *NA* | No record in WR2005 |
|  | *Lycalopex culpaeus* | *Lycalopex culpaeus* |  |
|  | *Lycalopex fulvipes* | *Lycalopex fulvipes* |  |
|  | *Lycalopex griseus* | *Lycalopex griseus* |  |
|  | *Lycalopex gymnocercus* | *Lycalopex gymnocercus* |  |
|  | *Lycalopex sechurae* | *Lycalopex sechurae* |  |
|  | *Lycalopex vetulus* | *Lycalopex vetulus* |  |
|  | *Lycaon pictus* | *Lycaon pictus* |  |
|  | *Nyctereutes procyonoides* | *Nyctereutes procyonoides* |  |
|  | *Otocyon megalotis* | *Otocyon megalotis* |  |
|  | *Speothos venaticus* | *Speothos venaticus* |  |
|  | *Urocyon cinereoargenteus* | *Urocyon cinereoargenteus* |  |
|  | *Urocyon littoralis* | *Urocyon littoralis* |  |
|  | *Vulpes bengalensis* | *Vulpes bengalensis* |  |
|  | *Vulpes cana* | *Vulpes cana* |  |
|  | *Vulpes chama* | *Vulpes chama* |  |
|  | *Vulpes corsac* | *Vulpes corsac* |  |
|  | *Vulpes ferrilata* | *Vulpes ferrilata* |  |
|  | *Vulpes lagopus* | *Vulpes lagopus* |  |
|  | *Vulpes macrotis* | *Vulpes macrotis* |  |
|  | *Vulpes pallida* | *Vulpes pallida* |  |
|  | *Vulpes rueppellii* | *Vulpes rueppellii* |  |
|  | *Vulpes velox* | *Vulpes velox* |  |
|  | *Vulpes vulpes* | *Vulpes vulpes* |  |
|  | *Vulpes zerda* | *Vulpes zerda* |  |
| *FELIDAE* | *Acinonyx jubatus* | *Acinonyx jubatus* |  |
|  | *Caracal aurata* | *Profelis aurata* |  |
|  | *Caracal caracal* | *Caracal caracal* |  |
|  | *Catopuma badia* | *Catopuma badia* |  |
|  | *Catopuma temminckii* | *Catopuma temminckii* |  |
|  | *Felis bieti* | *Felis bieti* |  |
|  | *Felis chaus* | *Felis chaus* |  |
|  | *Felis margarita* | *Felis margarita* |  |
|  | *Felis nigripes* | *Felis nigripes* |  |
|  | *Felis silvestris* | *Felis silvestris* |  |
|  | *Herpailurus yagouaroundi* | *Puma yagouaroundi* | Assigned to genus Puma in WR2005 |
|  | *Leopardus colocolo* | *Leopardus colocolo* |  |
|  | *Leopardus geoffroyi* | *Leopardus geoffroyi* |  |
|  | *Leopardus guigna* | *Leopardus guigna* |  |
|  | *Leopardus guttulus* | *NA* | Considered a sub-species of *Leopardus tigrinus* in WR2005 |
|  | *Leopardus jacobita* | *Leopardus jacobita* |  |
|  | *Leopardus pardalis* | *Leopardus pardalis* |  |
|  | *Leopardus tigrinus* | *Leopardus tigrinus* |  |
|  | *Leopardus wiedii* | *Leopardus wiedii* |  |
|  | *Leptailurus serval* | *Leptailurus serval* |  |
|  | *Lynx canadensis* | *Lynx canadensis* |  |
|  | *Lynx lynx* | *Lynx lynx* |  |
|  | *Lynx pardinus* | *Lynx pardinus* |  |
|  | *Lynx rufus* | *Lynx rufus* |  |
|  | *Neofelis diardi* | *NA* | Considered a sub-species of *Neofelis nebulosi* in WR2005 |
|  | *Neofelis nebulosa* | *Neofelis nebulosa* |  |
|  | *Otocolobus manul* | *Felis manul* | Assigned to genus Felis in WR2005 |
|  | *Panthera leo* | *Panthera leo* |  |
|  | *Panthera onca* | *Panthera onca* |  |
|  | *Panthera pardus* | *Panthera pardus* |  |
|  | *Panthera tigris* | *Panthera tigris* |  |
|  | *Panthera uncia* | *Uncia uncia* | Assigned to genus Uncia in WR2005 |
|  | *Pardofelis marmorata* | *Pardofelis marmorata* |  |
|  | *Prionailurus bengalensis* | *Prionailurus bengalensis* |  |
|  | *Prionailurus planiceps* | *Prionailurus planiceps* |  |
|  | *Prionailurus rubiginosus* | *Prionailurus rubiginosus* |  |
|  | *Prionailurus viverrinus* | *Prionailurus viverrinus* |  |
|  | *Puma concolor* | *Puma concolor* |  |
|  | *NA* | *Felis catus* | Species not considered by IUCN |
|  | *NA* | *Leopardus braccatus* | Species not considered by IUCN |
|  | *NA* | *Leopardus pajeros* | Species not considered by IUCN |
|  | *NA* | *Prionailurus iriomotensis* | Species not considered by IUCN |
| *HYAENIDAE* | *Crocuta crocuta* | *Crocuta crocuta* |  |
|  | *Hyaena hyaena* | *Hyaena hyaena* |  |
|  | *Parahyaena brunnea* | *Hyaena brunnea* | Assigned to genus Hyaena in WR2005 |
|  | *Proteles cristata* | *Proteles cristata* |  |
| *URSIDAE* | *Ailuropoda melanoleuca* | *Ailuropoda melanoleuca* |  |
|  | *Helarctos malayanus* | *Helarctos malayanus* |  |
|  | *Melursus ursinus* | *Melursus ursinus* |  |
|  | *Tremarctos ornatus* | *Tremarctos ornatus* |  |
|  | *Ursus americanus* | *Ursus americanus* |  |
|  | *Ursus arctos* | *Ursus arctos* |  |
|  | *Ursus maritimus* | *Ursus maritimus* |  |
|  | *Ursus thibetanus* | *Ursus thibetanus* |  |

**Table S3** Reference table for captrends.csv ‘Singular_country’ and ‘Multiple_countries’ fields. Country names follow ISO 3166 country name and two-character code standards. If sources described the global population trend, we added one row titled ‘GLOBAL’

| **Two-character code** | **Country** |
| --- | --- |
| AF | Afghanistan |
| AX | Åland Islands |
| AL | Albania |
| DZ | Algeria |
| AS | American Samoa |
| AD | Andorra |
| AO | Angola |
| AI | Anguilla |
| AQ | Antarctica |
| AG | Antigua and Barbuda |
| AR | Argentina |
| AM | Armenia |
| AW | Aruba |
| AU | Australia |
| AT | Austria |
| AZ | Azerbaijan |
| BS | Bahamas |
| BH | Bahrain |
| BD | Bangladesh |
| BB | Barbados |
| BY | Belarus |
| BE | Belgium |
| BZ | Belize |
| BJ | Benin |
| BM | Bermuda |
| BT | Bhutan |
| BO | Bolivia, Plurinational State of |
| BQ | Bonaire, Sint Eustatius and Saba |
| BA | Bosnia and Herzegovina |
| BW | Botswana |
| BV | Bouvet Island |
| BR | Brazil |
| IO | British Indian Ocean Territory |
| BN | Brunei Darussalam |
| BG | Bulgaria |
| BF | Burkina Faso |
| BI | Burundi |
| KH | Cambodia |
| CM | Cameroon |
| CA | Canada |
| CV | Cape Verde |
| KY | Cayman Islands |
| CF | Central African Republic |
| TD | Chad |
| CL | Chile |
| CN | China |
| CX | Christmas Island |
| CC | Cocos (Keeling) Islands |
| CO | Colombia |
| KM | Comoros |
| CG | Congo |
| CD | Congo, The Democratic Republic of the |
| CK | Cook Islands |
| CR | Costa Rica |
| CI | Côte D'Ivoire |
| HR | Croatia |
| CU | Cuba |
| CW | Curaçao |
| CY | Cyprus |
| CZ | Czech Republic |
| DK | Denmark |
| DJ | Djibouti |
| DM | Dominica |
| DO | Dominican Republic |
| EC | Ecuador |
| EG | Egypt |
| SV | El Salvador |
| GQ | Equatorial Guinea |
| ER | Eritrea |
| EE | Estonia |
| ET | Ethiopia |
| FK | Falkland Islands (Malvinas) |
| FO | Faroe Islands |
| FJ | Fiji |
| FI | Finland |
| FR | France |
| GF | French Guiana |
| PF | French Polynesia |
| TF | French Southern Territories |
| GA | Gabon |
| GM | Gambia |
| GE | Georgia |
| DE | Germany |
| GH | Ghana |
| GI | Gibraltar |
| GLOBAL | GLOBAL |
| GR | Greece |
| GL | Greenland |
| GD | Grenada |
| GP | Guadeloupe |
| GU | Guam |
| GT | Guatemala |
| GG | Guernsey |
| GN | Guinea |
| GW | Guinea-Bissau |
| GY | Guyana |
| HT | Haiti |
| HM | Heard Island and McDonald Islands |
| VA | Holy See (Vatican City State) |
| HN | Honduras |
| HK | Hong Kong |
| HU | Hungary |
| IS | Iceland |
| IN | India |
| ID | Indonesia |
| IR | Iran, Islamic Republic of |
| IQ | Iraq |
| IE | Ireland |
| IM | Isle of Man |
| IL | Israel |
| IT | Italy |
| JM | Jamaica |
| JP | Japan |
| JE | Jersey |
| JO | Jordan |
| KZ | Kazakhstan |
| KE | Kenya |
| KI | Kiribati |
| KP | Korea, Democratic People's Republic of |
| KR | Korea, Republic of |
| KW | Kuwait |
| KG | Kyrgyzstan |
| LA | Lao People's Democratic Republic |
| LV | Latvia |
| LB | Lebanon |
| LS | Lesotho |
| LR | Liberia |
| LY | Libya |
| LI | Liechtenstein |
| LT | Lithuania |
| LU | Luxembourg |
| MO | Macao |
| MK | Macedonia, The former Yugoslav Republic of |
| MG | Madagascar |
| MW | Malawi |
| MY | Malaysia |
| MV | Maldives |
| ML | Mali |
| MT | Malta |
| MH | Marshall Islands |
| MQ | Martinique |
| MR | Mauritania |
| MU | Mauritius |
| YT | Mayotte |
| MX | Mexico |
| FM | Micronesia, Federated States of |
| MD | Moldova, Republic of |
| MC | Monaco |
| MN | Mongolia |
| ME | Montenegro |
| MS | Montserrat |
| MA | Morocco |
| MZ | Mozambique |
| MM | Myanmar |
| NA | Namibia |
| NR | Nauru |
| NP | Nepal |
| NL | Netherlands |
| NC | New Caledonia |
| NZ | New Zealand |
| NI | Nicaragua |
| NE | Niger |
| NG | Nigeria |
| NU | Niue |
| NF | Norfolk Island |
| MP | Northern Mariana Islands |
| NO | Norway |
| OM | Oman |
| PK | Pakistan |
| PW | Palau |
| PS | Palestine, State of |
| PA | Panama |
| PG | Papua New Guinea |
| PY | Paraguay |
| PE | Peru |
| PH | Philippines |
| PN | Pitcairn |
| PL | Poland |
| PT | Portugal |
| PR | Puerto Rico |
| QA | Qatar |
| RE | Reunion |
| RO | Romania |
| RU | Russian Federation |
| RW | Rwanda |
| BL | Saint Barthélemy |
| SH | Saint Helena, Ascension and Tristan Da Cunha |
| KN | Saint Kitts and Nevis |
| LC | Saint Lucia |
| MF | Saint Martin (French Part) |
| PM | Saint Pierre and Miquelon |
| VC | Saint Vincent and the Grenadines |
| WS | Samoa |
| SM | San Marino |
| ST | Sao Tome and Principe |
| SA | Saudi Arabia |
| SN | Senegal |
| RS | Serbia |
| SC | Seychelles |
| SL | Sierra Leone |
| SG | Singapore |
| SX | Sint Maarten (Dutch Part) |
| SK | Slovakia |
| SI | Slovenia |
| SB | Solomon Islands |
| SO | Somalia |
| ZA | South Africa |
| GS | South Georgia and the South Sandwich Islands |
| SS | South Sudan |
| ES | Spain |
| LK | Sri Lanka |
| SD | Sudan |
| SR | Suriname |
| SJ | Svalbard and Jan Mayen |
| SZ | Swaziland |
| SE | Sweden |
| CH | Switzerland |
| SY | Syrian Arab Republic |
| TW | Taiwan, Province of China |
| TJ | Tajikistan |
| TZ | Tanzania, United Republic of |
| TH | Thailand |
| TL | Timor-Leste |
| TG | Togo |
| TK | Tokelau |
| TO | Tonga |
| TT | Trinidad and Tobago |
| TN | Tunisia |
| TR | Turkey |
| TM | Turkmenistan |
| TC | Turks and Caicos Islands |
| TV | Tuvalu |
| UG | Uganda |
| UA | Ukraine |
| AE | United Arab Emirates |
| GB | United Kingdom |
| US | United States |
| UM | United States Minor Outlying Islands |
| UY | Uruguay |
| UZ | Uzbekistan |
| VU | Vanuatu |
| VE | Venezuela |
| VN | Viet Nam |
| VG | Virgin Islands, British |
| VI | Virgin Islands, U.S. |
| WF | Wallis and Futuna |
| EH | Western Sahara |
| YE | Yemen |
| ZM | Zambia |
| ZW | Zimbabwe |

**Table S4.** Description of fields in the ts_abundance.csv table which provides the time series of population abundance estimates. The ‘Data type’ column describes the format of the data.

| **Field** | **Description** | **Data type** |
| --- | --- | --- |
| DataTableID | Unique numerical code for each source to match with table captrends.csv | Character |
| Value | Time-series value representing population abundance or density. | Numeric |
| Year | Time point of population abundance estimate (in years) | Numeric |

**Table S5.** Description of fields in the ts_change.csv table which provides the time series of population change estimates. The ‘Data type’ column describes the format of the data, for categorical fields the selection options are underlined and each options description is italicised.

| **Field** | **Description** | **Data type** |
| --- | --- | --- |
| DataTableID | Unique numerical code for each source to match with table captrends.csv | Character |
| Type_of_measure | Descriptor of the type of estimate in the time series, values presented in the ‘Value’ field. Categories:  *Lambda: estimate of the finite rate of population change between two time periods (represented by fields Year1 and Year2). 1 is stable*  *Percentage change: estimate of the percentage change in population size between two time periods (represented by fields Year1 and Year2). 100 is stable [formula = (N_t+1_/N_t_) * 100].* | Categorical |
| Value | Time-series value representing change in populations size. Interpreted alongside the Type_of_measure, Year1, and Year2 fields | Numeric |
| Year1 | Reference time-point (in years) e.g. date of first population estimate. | Numeric |
| Year2 | Change time-point (in years) e.g. date of second population estimate used to derive population change. | Numeric |

**Table S6.** Description of fields in the direction.csv table, which contains information on influences of the population trend, including whether the influence is positive or negative. This dataset uses existing classification schemes described in Table S6. The ‘Data type’ column describes the format of the data, for categorical fields the selection options are underlined and each options description is italicised.

| **Field** | **Description** | **Data type** |
| --- | --- | --- |
| DataTableID | Unique numerical code for each source to match with table captrends.csv | Character |
| Code | Amended threat or conservation action category described by the source as influencing the population trend. If an influencing factor could not be matched to a category, the driver of the trend, as described by the primary source, was entered as free text in the field ‘other_drivers_ot_trend’ in table captrends.csv | Categorical [calls on Table S7] |
| Direction | Binary descriptor of whether the factor was described by the source as potentially or actually having resulted, or being expected to result in a population increase (recorded as “Positive”) or in a population decline (recorded as “Negative”).  The degree of influence on the trend is informed by the Key_driver field. | Categorical |
| Measured | Descriptor of the evidence provided by the source to support the link between a named factor and changes in population trend. Categories:  *Not explained: sources mentioned potentially important factor but did not provide information on how it may affect population trend*  *Speculated: source speculated about a link between the factor and the population trend*  *Proxy-estimate: source provided some evidence for how the factor influenced the population trend*  *Quantified: source presented evidence that a factor has impacted the population trend* | Categorical |
| Key_driver | Descriptor of whether the factor was likely to be a strong driver of the observed population trend (recorded as “Positive”). Categories:  *No: Factor not considered an important driver of the trend according to primary source.*  *Yes: Factor considered an important driver of the trend according to primary source.*  *Unknown: Primary source did not describe impact of the factor, or described the impact as unknown.* | Categorical |
| Comment | Any additional notes regarding how the trend is influenced by the factor | Character |

**Table S7.** Reference table from the ‘Code’ field in the direction.csv file. When entering influences of population trends, the most detailed code possible was used. For example, if the source describes the trend as being influenced by small-scale fragmentation, 0.2.1 (Small-scale fragmentation) was selected. However, if the source describes the trend as being influenced by fragmentation, 0.2 (General fragmentation) was selected. Each ‘Code’ also falls within a higher level ‘Category’ which aggregates codes into broader groups. For each code the ‘Scheme description’ column provide the matching classifications in the IUCN Threats (scheme v3.2) and Conservation Actions (scheme v2.0) with the scheme specified in the ‘Scheme’ column as Threats or Conservation. Some threats and actions listed by sources were not well-matched to existing scheme categories, we created new Codes which are briefly described in the ‘Scheme description’ column and labelled as Added in the ‘Scheme’ column. Some categories from the IUCN schemes were not mentioned by the reviewed sources and were not used in the database. These are indicated with a ‘-‘ in the ‘Code’ column, and their scheme name is followed by an ‘X’.

| **Category** | **Code** | **Scheme description** | **Scheme** |
| --- | --- | --- | --- |
| **Habitat altered** | 0.1 (habitat altered - not targeted restoration) | 0.1 Habitat alteration but not restoration | Added |
|  | 0.1.1 (Small-scale habitat altered - not targeted restoration) | 0.1.1 Small scale alteration | Added |
|  | 0.1.2 (Large-scale habitat altered - not targeted restoration) | 0.1.2 Large-scale alteration | Added |
|  | 0.2 (General fragmentation) | 0.2 Fragmentation | Added |
|  | 0.2.1 (Small-scale fragmentation) | 0.2.1 Little fragmentation | Added |
|  | 0.2.2 (Large-scale fragmentation) | 0.2.2 Large-scale fragmentation | Added |
| **Residential & commercial development** | 1.1 (Habitat urbanised) | 1.1 Housing & urban areas | Threats |
|  | 1.2 (Habitat industrialised) | 1.2 Commercial & industrial areas | Threats |
|  | 1.3 (Habitat made available for recreation) | 1.3 Tourism & recreation areas | Threats |
| **Agriculture & aquaculture** | 2.1 (Habitat altered for farming) | 2.1 Annual & perennial non-timber crops | Threats |
|  | - | 2.1.1 Shifting agriculture | Threats X |
|  | 2.1.2 (Habitat altered for small-scale farming) | 2.1.2 Small-holder farming | Threats |
|  | 2.1.3 (Habitat altered for large-scale farming) | 2.1.3 Agro-industry farming | Threats |
|  | 2.2 (Habitat altered for plantations) | 2.2 Wood & pulp plantations | Threats |
|  | 2.2.1 (Habitat altered for small-scale plantations) | 2.2.1 Small-holder plantations | Threats |
|  | 2.2.2 (Habitat altered for large-scale plantations) | 2.2.2 Agro-industry plantations | Threats |
|  | - | 2.2.3 Scale unknown/Unrecorded | Threats X |
|  | 2.3 (Habitat altered for ranching) | 2.3 Livestock farming & ranching | Threats |
|  | 2.3.1 (Habitat altered for nomadic ranching) | 2.3.1 Nomadic grazing | Threats |
|  | 2.3.2 (Habitat altered for small-scale ranching) | 2.3.2 Small-holder grazing, ranching or farming | Threats |
|  | 2.3.3 (Habitat altered for large-scale ranching) | 2.3.3 Agro-industry grazing, ranching or farming | Threats |
|  | - | 2.3.4 Scale unknown/Unrecorded | Threats X |
|  | - | 2.4 Marine & freshwater aquaculture | Threats X |
|  | - | 2.4.1 Subsistence/artisanal aquaculture | Threats X |
|  | - | 2.4.2 Industrial aquaculture | Threats X |
|  | - | 2.4.3 Scale unknown/Unrecorded | Threats X |
| **Energy production & mining** | - | 3.1 Oil & gas drilling | Threats X |
|  | - | 3.2 Mining & quarrying | Threats X |
|  | - | 3.3 Renewable energy | Threats X |
| **Transportation & service corridors** | 4.1 (Road & railroads generally) | 4.1 Roads & railroads | Threats |
|  | 4.1.1 (Developing roads & rails) | 4.1.1 Roads & railroads development | Added |
|  | 4.1.2 (Vehicle collisions) | 4.1.2 Roads & railroads vehicle collisions | Added |
|  | - | 4.2 Utility & service lines | Threats X |
|  | - | 4.3 Shipping lanes | Threats X |
|  | - | 4.4 Flight paths | Threats X |
| **Human intrusions & disturbance** | 6.1 (Disturbance from recreational activities) | 6.1 Recreational activities | Threats |
|  | 6.2 (Disturbance from war) | 6.2 War, civil unrest & military exercises | Threats |
|  | 6.3 (Disturbance from people working) | 6.3 Work & other activities | Threats |
|  | - | 6.4 Other disturbance | Threats X |
| **Natural system modifications** | 7.1 (System altered by excess fire) | 7.1 Fire & fire suppression | Threats |
|  | 7.1.1 (System altered by fire shortage) | 7.1.1 Increase in fire frequency/intensity | Threats |
|  | - | 7.1.2 Suppression in fire frequency/intensity | Threats X |
|  | - | 7.1.3 Trend unknown/Unrecorded | Threats X |
|  | 7.2 (System altered by water shortage/dams) | 7.2 Dams & water management/use | Threats |
|  | 7.3 (System altered by ecosystem modifications) | 7.3 Other ecosystem modifications | Threats |
| **Invasive & other problematic species, genes & diseases** | 8.1 (Population effected by invasive disease) | 8.1 Invasive non-native/alien species/diseases | Threats |
|  | - | 8.1.1 Unspecified species | Threats X |
|  | - | 8.1.2 Named species | Threats X |
|  | 8.2 (Population effected by native disease) | 8.2 Problematic native species/diseases | Threats |
|  | - | 8.2.1 Unspecified species | Threats X |
|  | - | 8.2.2 Named species | Threats X |
|  | 8.3 (Population effected by introduced genes) | 8.3 Introduced genetic material | Threats |
|  | - | 8.4 Problematic species/diseases of unknown origin | Threats X |
|  | - | 8.4.1 Unspecified species | Threats X |
|  | - | 8.4.2 Named species | Threats X |
|  | - | 8.5 Viral/prion-induced diseases | Threats X |
|  | - | 8.5.1 Unspecified species | Threats X |
|  | - | 8.5.2 Named species | Threats X |
|  | 8.6 (Population effected by unknown disease) | 8.6 Diseases of unknown cause | Threats |
|  | - | 8.7 General disease | Threats X |
| **Pollution** | 9.1 (Population/Habitat effected by domestic waste) | 9.1 Domestic & urban waste water | Threats |
|  | 9.2 (Population/Habitat effected by industrial waste) | 9.2 Industrial & military effluents | Threats |
|  | - | 9.3 Agricultural & forestry effluents | Threats X |
|  | - | 9.4 Garbage & solid waste | Threats X |
|  | - | 9.5 Air-borne pollutants | Threats X |
|  | - | 9.6 Excess energy | Threats X |
|  | 9.6.1 (Population/Habitat effected by light pollution) | 9.6.1 Light pollution | Threats |
|  | 9.6.2 (Population/Habitat effected by thermal pollution) | 9.6.2 Thermal pollution | Threats |
|  | 9.6.3 (Population/Habitat effected by noise pollution) | 9.6.3 Noise pollution | Threats |
|  | - | 9.6.4 Type unknown/unrecorded | Threats X |
| **Geological events** | 10.1 (Population/Habitat effected by volcanoes) | 10.1 Volcanoes | Threats |
|  | 10.2 (Population/Habitat effected by earthquakes/tsunamis) | 10.2 Earthquakes/tsunamis | Threats |
|  | 10.3 (Population/Habitat effected by avalanches/landslides) | 10.3 Avalanches/landslides | Threats |
| **Climate change & severe weather** | 11.1 (Habitat shifts from climate change) | 11.1 Habitat shifting & alteration | Threats |
|  | 11.2 (Population/Habitat effected by drought) | 11.2 Droughts | Threats |
|  | 11.3 (Population/Habitat effected by temperature extremes) | 11.3 Temperature extremes | Threats |
|  | 11.4 (Population/Habitat effected by storms/flooding) | 11.4 Storms & flooding | Threats |
|  | 11.5 (Population/Habitat effected by unspecified climate change) | 11.5 Other impacts | Threats |
| **Biological resource use – adapted from section 5 of threats v3.2 to make all actions legal** | 13.1 (General legal hunting) | 5.1 Hunting & collecting terrestrial animals | Threats |
|  | 13.1.1 (Legal hunting of carnivore) | 5.1.1 Intentional use (species being assessed is the target) | Threats |
|  | 13.1.2 (Indirect effect from legal hunting) | 5.1.2 Unintentional effects (species being assessed is not the target) | Threats |
|  | 13.1.3 (Legal persecution/control of carnivore) | 5.1.3 Persecution/control | Threats |
|  | - | 5.1.4 Motivation unknown/Unrecorded | Threats X |
|  | 13.2 (Indirect effect of gathering plants) | 5.2 Gathering terrestrial plants | Threats |
|  | - | 5.2.1 Intentional use (species being assessed is the target) | Threats X |
|  | - | 5.2.2 Unintentional effects (species being assessed is not the target) | Threats X |
|  | - | 5.2.3 Persecution/control | Threats X |
|  | - | 5.2.4 Motivation unknown/Unrecorded | Threats X |
|  | - | 5.3 Logging & wood harvesting | Threats X |
|  | - | 5.3.1 Intentional use: subsistence/small scale (species being assessed is the target [harvest] | Threats X |
|  | - | 5.3.2 Intentional use: large scale (species being assessed is the target)[harvest] | Threats X |
|  | - | 5.3.3 Unintentional effects: subsistence/small scale (species being assessed is not the target)[harvest] | Threats X |
|  | - | 5.3.4 Unintentional effects: large scale (species being assessed is not the target)[harvest] | Threats X |
|  | - | 5.3.5 Motivation unknown/Unrecorded | Threats X |
|  | 13.4 (Indirect effect of fishing) | 5.4 Fishing & harvesting aquatic resources | Threats |
|  | - | 5.4.1 Intentional use: subsistence/small scale (species being assessed is the target)[harvest] | Threats X |
|  | - | 5.4.2 Intentional use: large scale (species being assessed is the target)[harvest] | Threats X |
|  | 13.4.3 (Indirect effect of small-scale fishing) | 5.4.3 Unintentional effects: subsistence/small scale (species being assessed is not the target) [harvest] | Threats |
|  | 13.4.4 (Indirect effect of large-scale fishing) | 5.4.4 Unintentional effects: large scale (species being assessed is not the target) [harvest] | Threats |
|  | - | 5.4.5 Persecution/control | Threats X |
|  | - | 5.4.6 Motivation unknown/Unrecorded | Threats X |
|  | 13.5 (Legal poisoning of carnivore) | 5.5 Poisoning | Added |
|  | 13.5.1 (Legal targeting poison towards carnivore) | 5.5.1 Intentional use (species being assessed is the target) | Added |
|  | 13.5.2 (Legal indirect poison of carnivore) | 5.5.2 Unintentional effects (species being assessed is not the target) | Added |
| **Biological resource use – adapted from section 5 of threats v3.2 to make all actions illegal** | 14.1 (General illegal hunting/poaching) | 5.1 Hunting & collecting terrestrial animals | Threats |
|  | 14.1.1 (Illegal hunting of carnivore/poaching) | 5.1.1 Intentional use (species being assessed is the target) | Threats |
|  | 14.1.2 (Indirect effect from illegal hunting/poaching) | 5.1.2 Unintentional effects (species being assessed is not the target) | Threats |
|  | 14.1.3 (Illegal persecution/control of carnivore) | 5.1.3 Persecution/control | Threats |
|  | - | 5.1.4 Motivation unknown/Unrecorded | Threats X |
|  | 14.2 (Indirect effect of gathering plants) | 5.2 Gathering terrestrial plants | Threats |
|  | - | 5.2.1 Intentional use (species being assessed is the target) | Threats X |
|  | - | 5.2.2 Unintentional effects (species being assessed is not the target) | Threats X |
|  | - | 5.2.3 Persecution/control | Threats X |
|  | - | 5.2.4 Motivation unknown/Unrecorded | Threats X |
|  | - | 5.3 Logging & wood harvesting | Threats X |
|  | - | 5.3.1 Intentional use: subsistence/small scale (species being assessed is the target [harvest] | Threats X |
|  | - | 5.3.2 Intentional use: large scale (species being assessed is the target)[harvest] | Threats X |
|  | - | 5.3.3 Unintentional effects: subsistence/small scale (species being assessed is not the target)[harvest] | Threats X |
|  | - | 5.3.4 Unintentional effects: large scale (species being assessed is not the target)[harvest] | Threats X |
|  | - | 5.3.5 Motivation unknown/Unrecorded | Threats X |
|  | 14.4 (Indirect effect of illegal fishing) | 5.4 Fishing & harvesting aquatic resources | Threats |
|  | - | 5.4.1 Intentional use: subsistence/small scale (species being assessed is the target)[harvest] | Threats X |
|  | - | 5.4.2 Intentional use: large scale (species being assessed is the target)[harvest] | Threats X |
|  | 14.4.3 (Indirect effect of small-scale illegal fishing) | 5.4.3 Unintentional effects: subsistence/small scale (species being assessed is not the target) [harvest] | Threats |
|  | 14.4.4 (Indirect effect of large-scale illegal fishing) | 5.4.4 Unintentional effects: large scale (species being assessed is not the target) [harvest] | Threats |
|  | - | 5.4.5 Persecution/control | Threats X |
|  | - | 5.4.6 Motivation unknown/Unrecorded | Threats X |
|  | 14.5 (Illegal poisoning of carnivore) | 5.5 Poisoning | Added |
|  | 14.5.1 (Illegal targeting poison towards carnivore) | 5.5.1 Intentional use (species being assessed is the target) | Added |
|  | 14.5.2 (Illegal indirect poison of carnivore) | 5.5.2 Unintentional effects (species being assessed is not the target) | Added |
| **Biological population drivers** | 15.1 (General competition) | 15.1 Competition | Added |
|  | 15.1.1 (Low competition - incomplete guild) | 15.1.1 Low inter-specific competition – incomplete carnivore guild | Added |
|  | 15.1.2 (Low competition - no reason) | 15.1.2 Low competition (reason not described) | Added |
|  | 15.1.3 (High competition - within guild) | 15.1.3 High inter-specific competition within guild | Added |
|  | 15.1.5 (High competition - no reason) | 15.1.5 High competition (reason not described) | Added |
|  | 15.1.6 (Low prey availability) | 15.1.6 Low prey availability | Added |
|  | 15.1.7 (High prey availability) | 15.1.7 High prey availability | Added |
|  | 15.1.8 (Competition from invasive species) | 15.1.8 Invasive non-native/alien species/diseases | Added |
|  | 15.2 (Carnivore predated) | 15.2 Predation | Added |
|  | 15.2.1 (Low predation risk - unbalanced guild ) | 15.2.1 Low predation risk – unbalanced guild | Added |
|  | 15.2.2 (Low predation risk - no reason) | 15.2.2 Low predation risk (reason not described | Added |
|  | 15.2.3 (High predation risk - unbalanced guild) | 15.2.3 High predation risk – unbalanced guild | Added |
|  | 15.2.4 (High predation risk - no reason) | 15.2.4 High predation risk (reason not described) | Added |
|  | 15.2.5 (Predation from invasive species) | 15.2.5 Invasive effects predation | Added |
|  | 15.3 (Below Minimum Viable Population) | 15.3 Population at minimum level | Added |
|  | 15.4 (High immigration/emigration) | 15.4 Population open | Added |
|  | 15.4.1 (High emigration) | 15.4.1 Emigration out of population | Added |
|  | 15.4.2 (High immigration) | 15.4.2 Immigration into population present | Added |
|  | 15.4.3 (Population expanding/recolonising areas) | 15.4.3 Range expansion/natural recolonization. | Added |
|  | 15.5 (Population closed/isolated) | 15.5 Population closed | Added |
|  | 15.5.1 (Low connectivity in population) | 15.5.1 Low connectivity | Added |
|  | 15.5.2 (Low connectivity - Inbreeding possible) | 15.5.2 Low connectivity – inbreeding possible | Added |
|  | 15.5.3 (Low connectivity - Inbreeding present) | 15.5.3 Low connectivity – inbreeding present | Added |
|  | 15.5.4 (No connectivity in population) | 15.5.4 No connectivity | Added |
|  | 15.5.5 (No connectivity - Inbreeding possible) | 15.5.5 No connectivity – inbreeding possible | Added |
|  | 15.5.6 (No connectivity - Inbreeding present) | 15.5.6 No connectivity – inbreeding present | Added |
|  | 15.5.7 (Unspecified genetic threat) | 15.5.7 Unspecified genetic threat | Added |
| **Land/water protection** | 1.1 (Protected area) | 1.1 Site/area protection | Conservation |
|  | 1.2 (Protected habitat) | 1.2 Resource & habitat protection | Conservation |
|  | 1.3 (Habitat developed over - considering sustainability) | 1.3 Resource & habitat protection | Conservation |
| **Land/water management** | 2.1 (Site managed) | 2.1 Site/area management | Conservation |
|  | 2.2 (Control of problematic species) | 2.2 Invasive/problematic species control | Conservation |
|  | 2.3 (Habitat restoration) | 2.3 Habitat & natural process restoration | Conservation |
| **Species management** | 3.1 (Species managed) | 3.1 Species management | Conservation |
|  | 3.1.1 (Harvest managed) | 3.1.1 Harvest management | Conservation |
|  | 3.1.2 (Trade managed) | 3.1.2 Trade management | Conservation |
|  | 3.1.3 (Population growth managed - culling) | 3.1.3 Limiting population growth | Conservation |
|  | 3.2 (Action to enable population recovery) | 3.2 Species recovery | Conservation |
|  | 3.3 (Re-introduction) | 3.3 Species re-introduction | Conservation |
|  | - | 3.3.1 Reintroduction | Conservation X |
|  | 3.3.2 (Benign introduction) | 3.3.2 Benign introduction | Conservation |
|  | 3.4 (Ex-situ - captive breeding/artificial propagation) | 3.4 Ex-situ conservation | Conservation |
|  | - | 3.4.1 Captive breeding/artificial propagation | Conservation X |
|  | - | 3.4.2 Genome resource bank | Conservation X |
| **Education & awareness** | 4.1 (Formal education) | 4.1 Formal education | Conservation |
|  | 4.2 (Train practitioners) | 4.2 Training | Conservation |
|  | 4.3 (Educate public) | 4.3 Awareness & communications | Conservation |
| **Law & policy** | 5.1 (General protective legislation) | 5.1 Legislation | Conservation |
|  | 5.1.1 (International legislation) | 5.1.1 International level | Conservation |
|  | 5.1.2 (National legislation) | 5.1.2 National level | Conservation |
|  | 5.1.3 (Regional legislation) | 5.1.3 Sub-national level | Conservation |
|  | - | 5.1.4 Scale unspecified | Conservation X |
|  | 5.2 (Protective policy) | 5.2 Policies and regulations | Conservation |
|  | - | 5.3 Private Sector Standards & Codes | Conservation X |
|  | 5.4 (Enforcing general policy/legislation) | 5.4 Compliance and enforcement | Conservation |
|  | 5.4.1 (Enforcing international policy/legislation) | 5.4.1 International level | Conservation |
|  | 5.4.2 (Enforcing National policy/legislation) | 5.4.2 National level | Conservation |
|  | 5.4.3 (Enforcing Regional policy/legislation) | 5.4.3 Sub-national level | Conservation |
|  | - | 5.4.4 Scale unspecified | Conservation X |
| **Livelihood, economic & other incentives** | 6.1 (Communities livelihood linked to species success)) | 6.1 Linked enterprises & livelihood alternatives | Conservation |
|  | 6.2 (Substitute carnivore for sustainable alternative) | 6.2 Substitution | Conservation |
|  | 6.3 (Carnivore market managed e.g. hunting levy) | 6.3 Market forces | Conservation |
|  | 6.4 (Compensation schemes) | 6.4 Conservation payments | Conservation |
|  | 6.5 (Utilise spiritual/religious connections for management) | 6.5 Non-monetary values | Conservation |

**Table S8.** Description of fields in the sources.csv table, which contains information on all reviewed sources. The ‘Data type’ column describes the format of the data, for categorical fields the selection options are underlined and each options description is italicised.

| **Field** | **Description** | **Data type** |
| --- | --- | --- |
| Year | Year of source publication | Numeric |
| Title | Title of source | Character |
| Citation_key | Unique alphanumerical identifier for each source which corresponds to captrends.csv | Character |
| Category | Category describing how the source was processed. Categories:  *Read – Data available: Population trend information was available within the source and extracted*  *Read – No Data available: Population trend information was unavailable within the source*  *NA: Source could not be accessed to assess if trend information was available.* | Categorical |
| From_Syst/Unsyst_search | Descriptor of how the source was found. Categories:  *1: Source identified through the unstructured search*  *2: Source identified through the structured search*  *3: Source identified through other sources (e.g. when reading category 1 or 2 sources, which mentioned other population trend values)* | Categorical |
| Comment | Any additional information regarding the source e.g. why trend data was not extracted. | Character |

**Table S9.** Reproducibility checklist from (Grames & Elphick, 2020). ‘Yes’ indicates information is available to reproduce the methods. ‘Partly’ describes the information as patially available. ‘No’ indicates that information is not available, and we describe the reason for this where listed. ‘NA’ indicates the category is not relevant to our methods. We only scored the ‘systematic’ search in this table, acknowledging that the small proportion of additional (~10%) studies identified in our unstructured search are not easily reproducible.

|  | Yes/Partly/No |
| --- | --- |
| **Is the search strategy described in sufficient detail to be repeatable by an independent**  **researcher? Given the search strategy, would an independent researcher be able to obtain the same full set of articles retrieved in the original search?** |  |
| Full search string is provided for at least one database or an unambiguous method of  reconstructing the search string is described | Yes |
| Databases and gray literature sources searched are listed | No – Databases in Web of Knowledge and Scopus were not noted at the point of the search. |
| Years of database access are provided | Yes |
| Date of search or last accession date is provided | Yes |
| **Is the screening process described in sufficient detail to be repeatable by an independent**  **researcher? Given the results of the original search, would an independent researcher be able to repeat the process and end up with the same set of included studies?** | Yes |
| Inclusion and/or exclusion criteria are provided | Yes |
| Inclusion/exclusion criteria include operational definitions | Yes |
| The method of screening is described in sufficient detail that the decision making  process is transparent, or a screening protocol is provided, or reasons for exclusion  at the full text stage are given | Yes |
| **Is the assessment of primary study quality described in sufficient detail to be repeatable by an independent researcher? Given the list of included studies, would an independent researcher be able to re-assess their quality and arrive at the same appraisals as the review?** | Yes |
| A standard tool was used and referenced or the critical appraisal process was described with sufficient detail in the review itself | Yes |
| **Is the data coding and extraction described in sufficient detail to be repeatable by an**  **independent researcher? Given the list of included studies, would an independent researcher be able to reconstruct the data set used for inference?** | Yes |
| Variables coded are provided | Yes |
| Variables coded have operational definitions | Yes |
| If a quantitative analysis is performed, any transformations of the data (e.g. effect  size calculations, re-coding results of studies, etc.) are described in sufficient detail | NA |
| **Are the data synthesis methods adequately described? Given the same data set, would an**  **independent researcher arrive at the same results?** | NA |
| For narrative reviews, the methods of synthesizing results across studies are  described, or for quantitative synthesis, the methods or models are described in  sufficient detail that the analysis could be redone | NA |
